# *Sorghum bicolor* cultivars have divergent and dynamic gene regulatory networks that control the temporal expression of genes in stem tissue

**DOI:** 10.1101/2020.06.17.158048

**Authors:** Madara Hetti Arachchilage, John E. Mullet, Amy Marshall-Colon

**Affiliations:** Center for Advanced Bioenergy and Bioproducts Innovation BRC, University of Illinois Urbana-Champaign, Urbana, IL, USA 61801; Department of Biochemistry and Biophysics, Texas A&M University, College Station, TX, 77843 USA; Department of Plant Biology, University of Illinois Urbana-Champaign, 265 Morrill Hall, MC-116, 505 South Goodwin Ave., Urbana, IL 61801, USA

**Keywords:** Stem-specific genes, *BTx623*, *Sweet Della*, *Sorghum bicolor*, Tissue-specific gene regulatory network, Spatial-temporal gene regulation, Cis-regulatory elements, Single nucleotide variants, Structural variants

## Abstract

The genetic engineering of value-added traits such as the accumulation of bioproducts in high biomass C4 grass stems is one promising strategy to make plant-derived biofuels more economical for industrial use. A first step toward achieving this goal is to identify stem-specific promoters that can drive the expression of genes of interest with good temporal and spatial specificity. However, a comprehensive characterization of the spatial-temporal regulatory elements of stem tissue-specific promoters for C4 grasses has not been reported. Therefore, we performed an *in-silico* analysis on *Sorghum bicolor cv BTx623* transcriptomes from multiple tissues over development to identify stem-expressed genes. The analysis identified 10 genes that are “Always-On-Stem-Specific,” 59 genes that are “Temporally-Stem-Specific during early development,” and 21 genes that are “Temporally-Stem-Specific during late development.” Promoter analysis revealed common and/or unique cis-regulatory elements in promoters of genes within each of the three categories. Subsequent gene regulatory network (GRN) analysis revealed that different transcriptional regulatory programs are responsible for the temporal activation of the stem-expressed genes. The analysis of temporal stem GRNs between sweet (*cv Della*) and grain (*cv BTx623*) sorghum varieties revealed genetic variation that could influence the regulatory landscape. This study provides new insights about sorghum stem biology, and information for future genetic engineering efforts to fine-tune the spatial-temporal expression of transgenes in C4 grass stems.

## Introduction

High biomass grasses, such as sorghum (*Sorghum bicolor*), miscanthus (*Miscanthus x giganteus*), switchgrass (*Panicum virgatum*), and sugarcane (*Saccharum sp.*), offer an abundant and renewable source of biomass for biofuels production (McLaughlin and Kszos, 2005; Rooney *et al.*, 2007; Karp and Shield, 2008; Murray *et al.*, 2008; Waclawovsky *et al.*, 2010a; Byrt, Grof and Furbank, 2011; Sage and Zhu, 2011; Feltus and Vandenbrink, 2012; Olson *et al.*, 2012; Mullet *et al.*, 2014; Mathur *et al.*, 2017) providing a sustainable alternative to conventional fossil fuels (Demirbas, 2007; Demirbas and Demirbas, 2010; Kumar, Long and Singh, 2018; Kumar, 2020). Grasses with C4 photosynthesis can produce high biomass in marginal lands with low water and nitrogen inputs, leading to energy conservation (McLaughlin *et al.*, 2002; Heaton, Dohlemnn and Long, 2008; Varvel *et al.*, 2008). However, the cost associated with production of fuels and other bioproducts from plant biomass is relatively high, which has been a major constraint on the widespread adoption of grasses as sources of feedstock to produce biofuels or bioproducts (Hill *et al.*, 2006; Kumar, Long and Singh, 2018). Previous work showed that C4 grasses can be genetically engineered for increased fatty acid production, resulting in biomass with high oil content (Huang, Long and Singh, 2015; Huang *et al.*, 2016; Wang, 2016; Zale *et al.*, 2016; Beechey-Gradwell *et al.*, 2020; Mitchell *et al.*, 2020; Parajuli *et al.*, 2020), which increases their value as biofuels. Genetically engineering high-biomass grasses to accumulate valuable bioproducts that require minimal processing post-harvest is a desirable way forward, but has many challenges.

Seeds are the primary site of lipid synthesis and storage in plants (Harwood *et al.*, 1971; Dyer and Mullen, 2005; Lung and Weselake, 2006; Graham, 2008; Koçar and Civaş, 2013; Li *et al.*, 2016; Baud, 2018) in the form of triacylglycerols (TAGs) which have twice the energy density of carbohydrates and can be easily converted to other biofuel products. While oil-accumulating seeds have been a source of lipids for the biofuel industry, use of oil seeds or fruits for biofuels could negatively impact food supplies and prices. Therefore, genetic engineering of vegetative tissues to accumulate higher triacylglycerol (TAG) levels has drawn attention (Durrett, Benning and Ohlrogge, 2008; Qazi, Paranjpe and Bhargava, 2012; Napier *et al.*, 2014a; Kumar, 2020). Since C4 grasses can be grown on marginal lands not suitable for most food crops, the high biomass producing vegetative tissues of sorghum, miscanthus, and sugarcane are desirable targets for engineering efforts aimed at producing and/or storing lipids (Carlsson *et al.*, 2011; Sanjaya *et al.*, 2011; Weselake, 2016a; Lima *et al.*, 2017). For example, stem tissues engineered to divert energy from nonstructural carbohydrates into the lipid biosynthesis pathway through strategies such as increasing the supply of FAs, increasing TAG assembly activities, and blocking TAG breakdown pathways have resulted in higher amounts of TAGs accumulating in vegetative tissues (Papini-Terzi *et al.*, 2009; Waclawovsky *et al.*, 2010b; Sanjaya *et al.*, 2011; Qazi *et al.*, 2014; Sekhon *et al.*, 2016). Other proof-of-concept studies for increasing TAGs in vegetative tissues have been performed in *Arabidopsis thaliana, Brachypodium distachyon, Nicotiana benthamiana, Nicotiana tabacum*, sugarcane (*Saccharum* spp.), and *Sorghum bicolor* (Thelen and Ohlrogge, 2002; Fan, Yan and Xu, 2013; Vanhercke *et al.*, 2013, 2019; Yang *et al.*, 2015; Zale *et al.*, 2016; Mitchell *et al.*, 2020; Parajuli *et al.*, 2020; Xu *et al.*, 2020). For example, in engineered sugarcane, TAGs accumulated to an average of 8.0% of the dry weight of leaves and 4.3% of the dry weight of stems (Huang, Long and Singh, 2015; Zale *et al.*, 2016; Parajuli *et al.*, 2020). Another study in tobacco achieved TAGs accumulation up to 19% of the dry weight of the total biomass production by over-expressing the genes encoding WRINKLED1, DGAT, and oleosins (Vanhercke *et al.*, 2014; Zale *et al.*, 2016). However, many of these engineering efforts to increase TAG accumulation in immature vegetative tissues have resulted in negative impacts on plant growth as observed in sorghum, potato, tobacco, and Arabidopsis (Slocombe *et al.*, 2009; Feltus and Vandenbrink, 2012; Kelly *et al.*, 2013; Vanhercke *et al.*, 2014, 2019; Hofvander *et al.*, 2016; Liu *et al.*, 2017; Ramšak *et al.*, 2018; Xiaoyu Xu *et al.*, 2019; Xu *et al.*, 2020; Mitchell *et al.*, 2020). One hypothesis is that driving lipid accumulation under the control of tissue- and/or developmental-stage specific promoters, specifically those active during late development (Moyle and Birch, 2013; Mudge *et al.*, 2013), will have less of an impact on photosynthetic efficiencies and plant growth than constitutive overexpression of genes of interest.

Sink–source dynamics within the plant direct how much, where, and when carbohydrates are allocated, and determine final biomass of vegetative plant organs and seed. Knowledge of the spatial and temporal heterogeneity of carbon sinks among tissues and cell types in vegetative tissues during development could enable more informed genetic engineering (Schnyder, 1993; Hoffmann-Thoma *et al.*, 1996; Wang *et al.*, 2007; Sanjaya *et al.*, 2011; Slewinski, 2012; da Costa *et al.*, 2014; Sekhon *et al.*, 2016; Hennet *et al.*, 2020). For example, previous studies reported that sorghum, sugarcane, and wheat store excess carbohydrates in the form of soluble sugars in stem pith parenchyma, in close proximity to vascular bundles located within internode, and these stem carbohydrate reserves can be remobilized and partitioned to the developing grain during the reproductive phase (Hoffmann-Thoma *et al.*, 1996; Douglas *et al.*, 2002; Walsh, Sky and Brown, 2005; Ruuska *et al.*, 2006; Tarpley and Vietor, 2007; Rae *et al.*, 2009; Qazi, Paranjpe and Bhargava, 2012; Sekhon *et al.*, 2016; Kebrom, McKinley and Mullet, 2017; McKinley *et al.*, 2018). In such cases, tissue and developmental-stage specific promoters could be leveraged to divert the nonstructural carbohydrates to tissues engineered for elevated TAG production and accumulation, potentially avoiding the previously observed inhibition of photosynthesis and growth at early developmental stages. Many tissue-specific promoters have been successfully characterized and applied to targeted genetic engineering of maize, rice, soybean, tomato, potato, and tobacco (Deikman, Kline and Fischer, 1992; Trindade *et al.*, 2003; Cao *et al.*, 2007; Li *et al.*, 2013, 2019; Molla *et al.*, 2013; Dutt *et al.*, 2014; Napier *et al.*, 2014b, 2014a; Chen *et al.*, 2015; Wang *et al.*, 2017). Thus, increasing oil accumulation in stems of biofuel crops without impacting plant growth will require manipulation of developmental-stage specific and stem-specific promoters. Potential strategies include: a) converting sucrose to lipid in specific cells/tissues at stages when non-structural carbohydrates are highly abundant, and b) increasing stem and other vegetative tissue carbon sink strength, and sink storage capacity at late developmental stages (Slocombe *et al.*, 2009; Zhang *et al.*, 2010; P. Joshi and Nookaraju, 2012; Kim *et al.*, 2015; Weselake, 2016b; Xu and Shanklin, 2016; Shih, 2018).

Cis -regulatory elements in promoters upstream of a gene provide specific binding sites for their corresponding transcription factors (TFs), which regulate the transcription of gene expression. Studies in Arabidopsis and a few crop species have uncovered cis-regulatory elements that govern stem vascular tissue specific expression patterns, including phloem-specific, xylem-specific, and companion cell-specific promoter elements (***Table S1***) (Dutt *et al.*, 2014). However, a comprehensive characterization of the spatial-temporal regulatory elements of stem specific promoters for C4 grasses has not been reported but is necessary to understand the regulation of stem-expressed genes. Therefore, we performed an *in-silico* genome-wide analysis to identify stem-specific genes in Sorghum and the regulatory network that drives their spatial-temporal expression dynamics. We also examined the conservation/divergence of TF regulators of stem-specific genes between the two sorghum cultivars, sweet sorghum (*Sorghum bicolor cv* Della) and grain sorghum (*Sorghum bicolor cv* BTx623).

## Results and Discussion

We analyzed a gene expression atlas of sorghum tissues (McCormick *et al.*, 2018a) to identify genes specifically expressed in stem tissue over development. For this purpose, we used 38 expression profiles of five different *Sorghum bicolor* (*cv* BTx623) organs (stem, leaves, roots, peduncle, panicle, seeds) across five developmental stages (juvenile, vegetative, floral initiation, anthesis, grain maturity) to perform a tissue specificity analysis. We explored the gene regulatory networks (GRNs) for stem-expressed genes in an attempt to identify regulatory promoter elements that may contribute to the tissue-specific expression of these genes. The identification of such elements may help to uncover a stem-specific promoter that could be used as a molecular tool for the temporal and spatial expression of genes of interest that are not normally expressed in stem tissue. We performed further investigation to identify genetic variations in stem-preferred genes and their regulators, which cause large changes in the stem-specific GRNs between sweet sorghum (*Sorghum bicolor cv* Della) and grain sorghum (*Sorghum bicolor cv* BTx623).

### Tau index identifies stem-expressed genes over development

Several methods have been developed to determine tissue-specific gene expression (Huminiecki, Lloyd and Wolfe, 2003; Schug *et al.*, 2005; Yanai *et al.*, 2005; Smith *et al.*, 2006; Yu *et al.*, 2006; Xiao *et al.*, 2010; Julien *et al.*, 2012; Uhlen *et al.*, 2015; O’Hagan *et al.*, 2018). A recent benchmark study compared nine algorithms to determine tissue specificity and identified that the Tau (τ) index was the most robust analysis (Kryuchkova-Mostacci and Robinson-Rechavi, 2016). According to the tissue-specificity benchmark study, τ value analysis identified the most known tissue-specific genes in the human genome; including genes not detected by other algorithms, and avoided bias toward classifying most genes as “housekeeping.”

In this study, we used τ index to estimate the tissue-specificity of expressed genes at a given developmental stage. A summary of the number of tissue-specific genes found for each tissue type at each development stage is listed in ***Table S2***. A small portion of genes exhibited tissue-specific expression at each of the developmental stages (τ = 1) in stem, root, leaf, peduncle, panicle and seed tissues. There were fewer tissue-specific genes in stem tissue compared to leaf, root, and reproductive tissues. We observed a higher number of stem-specific genes during early stages compared to late stages (157, 133, 21, 21 and 44 stem-specific genes at juvenile, vegetative, floral, anthesis, and grain stages, respectively). Of the stem-specific genes identified at each developmental stage, some genes were always expressed in the stem throughout development, while some genes were transiently stem-specific over development. In this study, we classified three groups of stem-expressed genes: AOSS (Always-On-Stem-Specific), TSS-Early (Temporally-Stem-Specific during early development), and TSS-Late (Temporally-Stem-Specific during late development), in which there were 10 AOSS, 59 TSS-Early, and 21 TSS-Late genes (***Table S3***).

### Expression patterns of AOSS, TSS-Early, TSS-Late genes and their functional characterization

There were 10 genes in the AOSS group that had high stem-specificity (no significant expression at threshold of 5 TPM in non-stem tissues). Specifically, Sobic.006G024800 (an Andropogoneae family gene), Sobic.006G026300 (a Poaceae family gene) and Sobic.003G272000 (a Poales family gene) had high tissue specificity (τ = 1) and high expression (expression > 100 TPM) across the five developmental time points (***Figure 2, Figure S1***). The high stem-specificity value for these genes in *Sorghum cv BTx623* make them interesting candidates for further investigation. Other genes in the AOSS group had high stem-specificity at early developmental stages; however, during the later stages these genes were expressed at similar levels among vegetative stems, the developing peduncle, and panicles which also contain stem tissue (rachis, pedicles) (***Figure 2***).These genes include Sobic.001G106200 (KNAT1; TALE TF family gene with KNOX DNA binding domain), Sobic.001G106000 (KNAT1; TALE TF), Sobic.010G194600 (HSP90-like ATPase family protein), Sobic.006G147500 (light sensitive hypocotyls 6, LSH6), Sobic.002G419500 (HAD superfamily, subfamily IIIB acid phosphatase), Sobic.003G007200 (TIP4; tonoplast intrinsic protein), Sobic.007G008700 (ARM repeat superfamily protein, Poaceae specific family). KNAT1 is involved in meristem maintenance promoting cell proliferation and inhibiting differentiation and was previously shown to move between adjacent cells via intercellular pores formed by plasmodesmata in Arabidopsis (Kehr & Kragler, 2018). In Arabidopsis, *knat1* mutant plants had reduced formation of xylem fibers (Helizon *et al.*, 2014a).

**Figure 1:**
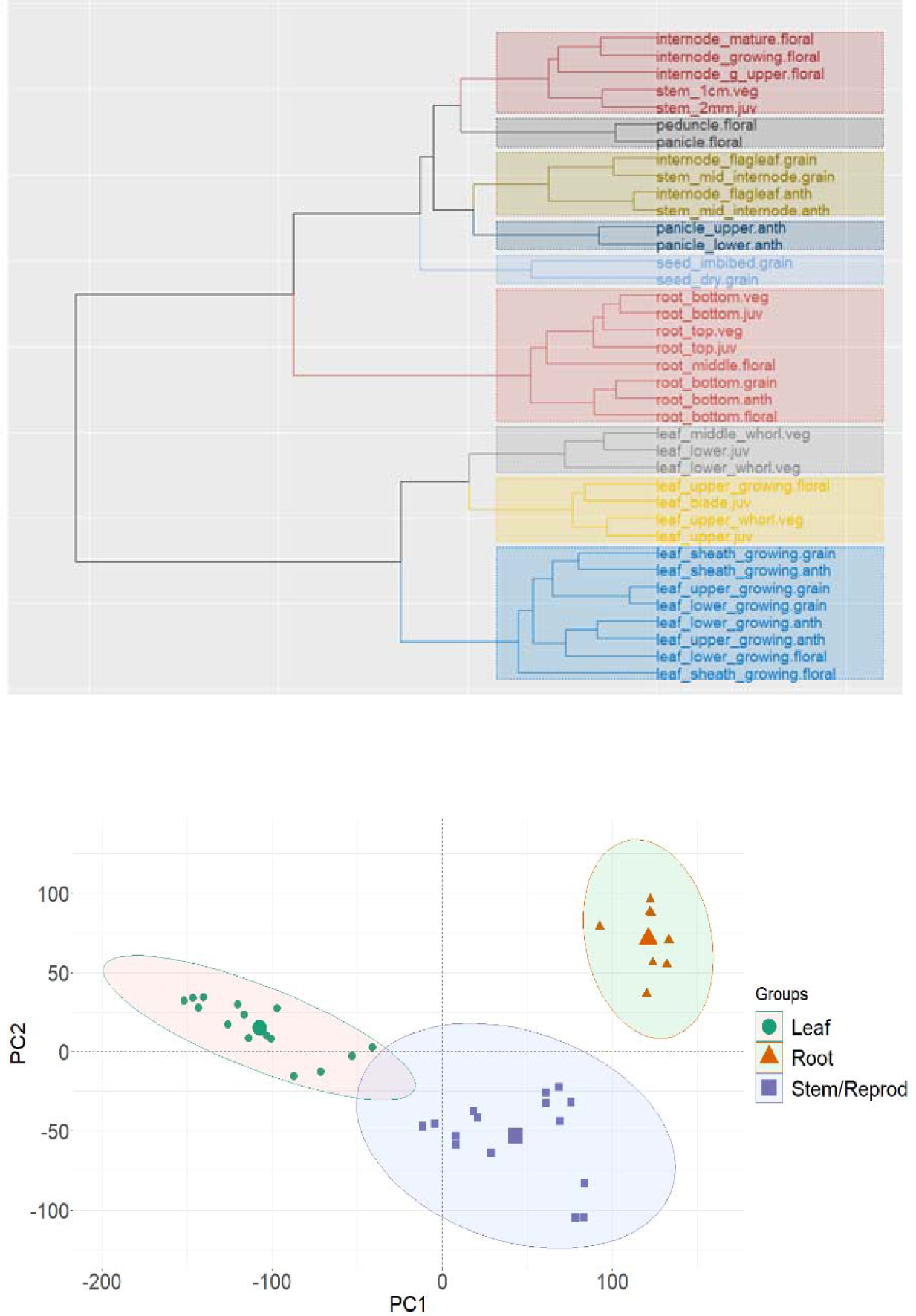
Clustering of tissues. A) Hierarchical Clustering analysis of the 38 tissue samples based on the expression of 23,674 genes (expressed in at least one sample, TPM ≥ 5). B) First two principal components of the spatial-temporal samples colored based on 4 major tissue types using expression values of all genes.

**Figure 2:**
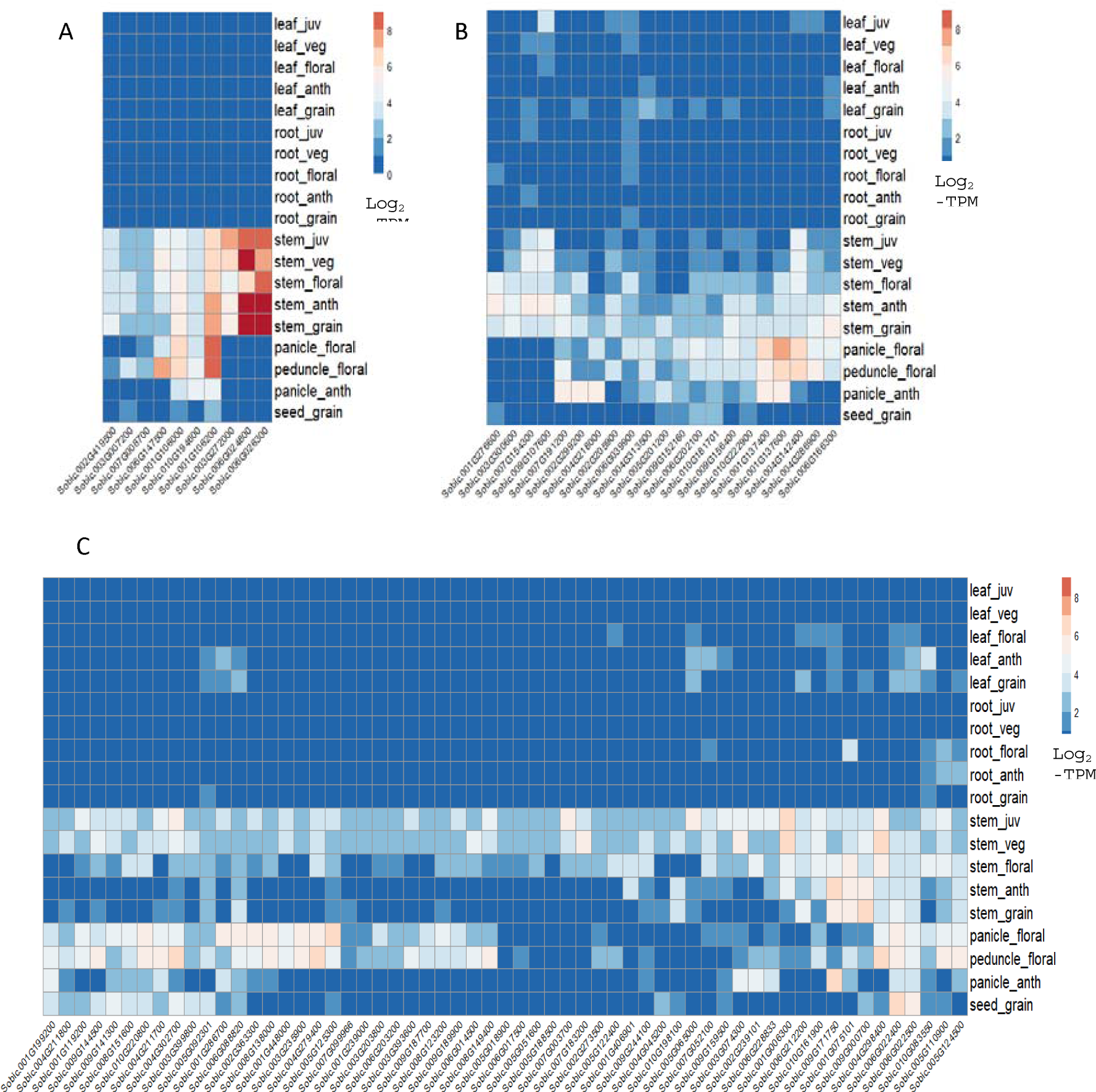
Global gene expression of stem-specific genes across stem and non-stem tissues through all developmental stages. A) Gene expression profile of 10 AOSS (Always-On-Stem-Specific) genes, B) Gene expression profile of 59 TSS-Early (Temporal stem-specificity during Early development), B) Gene expression profile of 21 TSS-Late (Temporal Stem-Specificity during Late development). Each row represents the mean gene expression values/ quantile normalized log2-TPM values. The mean expression values for each gene in each stage are represented by the intensity of color (blue representing no expression and orange representing maximum expression).

There were 59 TSSE genes that are “stem-preferred” rather than stem-specific, which are highly expressed in the stem during early development, but have no, low, or moderate expression in non-stem tissues (leaf, root, seed) during late development. The TSSE genes that have high stem specificity (τ = 1) during vegetative and juvenile stages and are not expressed during later stages (***Table S3***), including Sobic.001G239000, Sobic.005G018900 (PLC2; Phospholipase C2), Sobic.005G051600 (DUF247), Sobic.005G188500 (DUF94), Sobic.006G017500 (Disease resistance protein family/CC-NBS-LRR class), and Sobic.007G099966 (Andropogoneae specific family). Other TSSE genes also have high stem specificity (τ = 1) during vegetative and juvenile stages but have low to moderate expression in non-stem tissues during late development, thus their tissue specificity is more temporal. Many of the genes in this latter group have known stem-related functions related to early regulation of vascular tissue development and maintenance as listed in ***Table 2***. Two of these genes, YAB1 and CLAVATA3, are involved in CLAVATA (CLV)-WUSCHEL (WUS) pathway to regulate stem cell maintenance in shoot and floral meristems (Caño-Delgado, Lee and Demura, 2010; Ruonala, Ko and Helariutta, 2017). CLVA3 is expressed in the central domain of SAM and influences the relative positioning of the organ primordia by acting together with CLV1 and WUS (Ottoline Leyser and Furner, 1992), while YAB1 defines the primordia domain of the SAM periphery and promotes robust partitioning of the SAM by non-autonomously communicating with the meristem to regulate the expression of CLV3 and WUS. This coordination is essential for organized growth of the SAM (Goldshmidt *et al.*, 2008a; Stahle *et al.*, 2009a).

**Table 1:** The information of tissues sampled by development stage used in the transcriptome dataset used in this analysis from [McCormick et al., 2018].

| Tissue Type | Stage | Time of sample | Tissue Description | Sample ID |
| --- | --- | --- | --- | --- |
| Leaves | Juvenile | DAE 8 | Leaf blade of first emerged leaf | leaf.blade.juv |
|  |  |  | Lower portion (proximal) of third emerging leaf blade (in whorl) | leaf.lower.juv |
|  |  |  | Upper portion (distal) of third emerging leaf blade | leaf.upper.juv |
| Roots |  |  | Central section of roots | root_bottom.juv |
|  |  |  | Distal 1cm of root tips | root_top.juv |
| Stem |  |  | Entire stem (approximately 2mm in size) | stem_2mm.juv |
| Leaves | Vegetative | DAE 24 | Lower portion (proximal) of shoot, including sheaths, after removing first through fourth emerged leaves | leaf_lower_whorl.veg |
|  |  |  | Middle portion of shoot after removing first through fourth emerged leaves | leaf_middle_whorl.veg |
|  |  |  | Upper portion (distal) of shoot, including leaves emerged from whorl, after removing first through fourth emerged leaves | leaf_upper_whorl.veg |
| Roots |  |  | Central section of roots | root_bottom.veg |
|  |  |  | Distal 1cm of root tips | root_top.veg |
| Stem |  |  | Entire stem (approximately 1cm in size) | stem_1cm.veg |
| Leaves | Floral initiation | DAE 44 | Leaf sheath of last fully emerged (ligulated) leaf | leaf_sheath_growing.floral |
|  |  |  | Lower portion (proximal) of leaf blade of last fully emerged leaf (ligulated) | leaf_lower_growing.floral |
|  |  |  | Upper portion (proximal) of leaf blade of last fully emerged leaf (ligulated) | leaf_upper_growing.floral |
| Roots |  |  | Central section of roots | root_middle.floral |
|  |  |  | Distal 1cm of root tips | root_bottom.floral |
| Stem |  |  | Mature internode from middle of stem | internode_mature.floral |
|  |  |  | 2cm long growing internode near apex (further from apex than 1cm growing internode) | internode_growing.floral |
|  |  |  | 1cm long growing internode near apex (closer to apex than 2cm growing internode) | internode_growing_upper.floral |
| Reproductive Structures |  |  | Region of developing panicle in apex | panicle.floral |
|  |  |  | Region of developing peduncle in apex | peduncle.floral |
| Leaves | Anthesis | DAE 65 | Leaf sheath of leaf below flag leaf | leaf_sheath_growing.anth |
|  |  |  | Lower portion (proximal) of leaf blade of leaf below flag leaf | leaf_lower_growing.anth |
|  |  |  | Upper portion (distal) of leaf blade of leaf below flag leaf | leaf_upper_growing.anth |
| Roots |  |  | Central section of roots | root_bottom.anth |
| Stem |  |  | Mature internode from middle of stem | stem_mid_internode.anth |
|  |  |  | Mature internode immediately below peduncle | Internode_flagleaf.anth |
| Reproductive Structures |  |  | Lower (proximal) half of panicle | panicle_lower.anth |
|  |  |  | Upper (distal) half of panicle | panicle_upper.anth |
| Leaves | Grain maturity | DAE 96 | Leaf sheath of flag leaf | leaf_sheath_growing.grain |
|  |  |  | Lower portion (proximal) of flag leaf blade | leaf_lower_growing.grain |
| Roots |  |  | Central section of roots | root_bottom.grain |
| Stem |  |  | Mature internode from middle of stem | Stem_mid_internode.grain |
|  |  |  | Mature internode immediately below peduncle | Internode_flagleaf.grain |
| Reproductive Structures |  |  | Mature, dormant, dry seed | Seed_dry.grain |
|  |  |  | Developing seed, imbibed overnight | Seed_imbibed.grain |

**Table 2:**
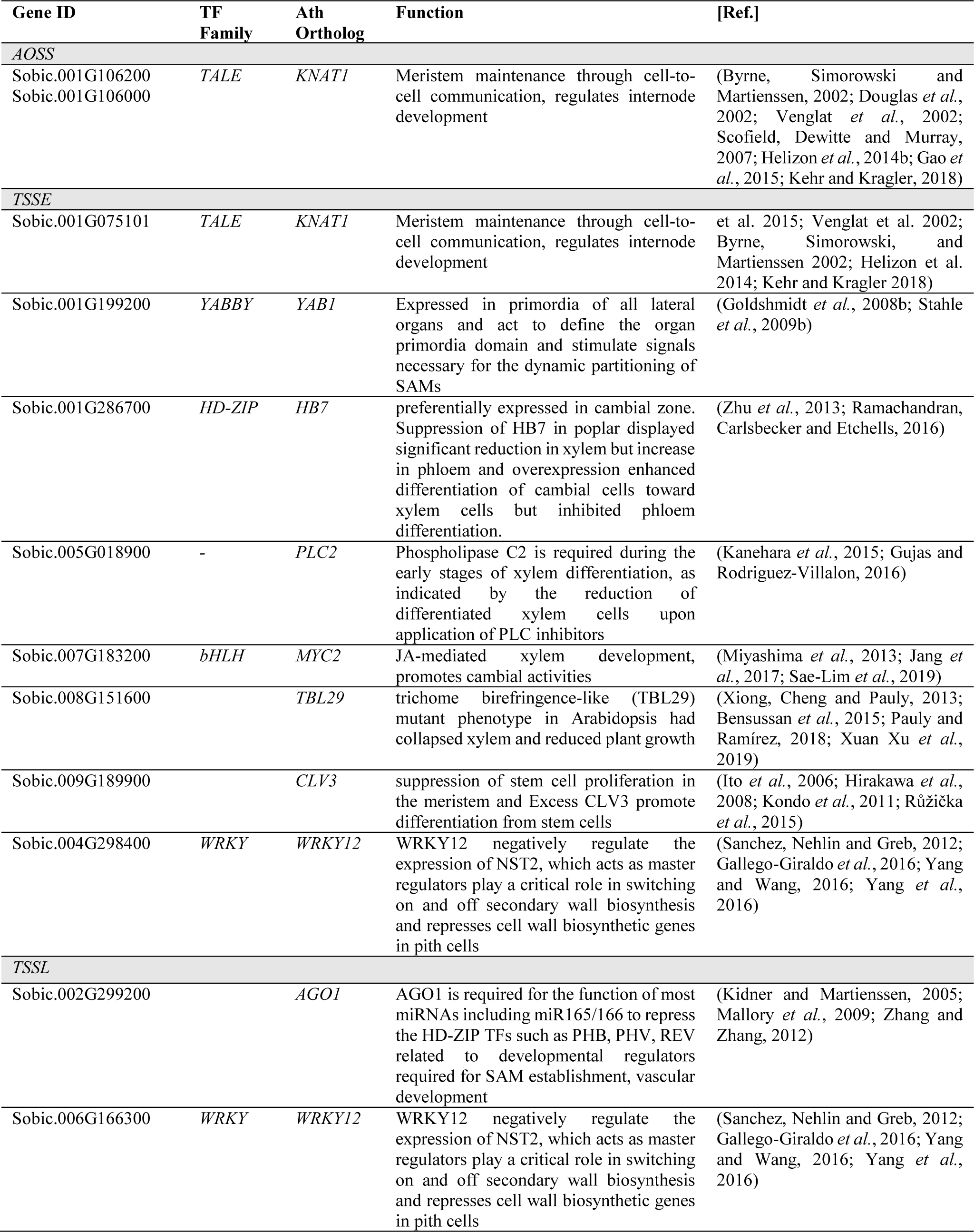
Stem-related cellular functions of identified stem-specific and stem-preferred genes.

There were 21 TSSL genes that, like the TSSE genes, are stem-preferred temporally during late development but have no, low, or moderate expression in non-stem tissues (leaf, root) during early development (***Table S3***). Two of these TSSL genes that have high stem specificity during floral, anthesis, and grain maturity stages, but are expressed in non-stem tissues during early development, include Sobic.002G299200 (AGO1) and Sobic.006G166300 (WRKY12). AGO1 is required for the function of most miRNAs, including miR165/166, to repress the HD-ZIP TFs such as PHB, PHV, REV related to the developmental regulators required for SAM establishment and vascular development in *Arabidopsis* (Kidner and Martienssen, 2005; Mallory *et al.*, 2009; Zhang and Zhang, 2012). WRKY12 was previously shown to negatively regulate the expression of NST2, which acts as a master regulator in switching on and off secondary cell wall biosynthesis and represses cell wall biosynthetic genes in pith cells in *Arabidopsis, Medicago sativa L.* (Sanchez, Nehlin and Greb, 2012; Gallego-Giraldo *et al.*, 2016; Yang and Wang, 2016; Yang *et al.*, 2016). Genes involved in secondary cell wall formation were down-regulated in Della post anthesis, possibly mediates in part through the action of WRKY12 (McKinley et al., 2016).

To elucidate the enriched functional categories of the identified stem-preferred genes, GO term analysis was performed using PlantTFDB for AOSS, TSSE and TSSL gene sets separately (***Figure 3***). AOSS and TSSE genes were enriched in biological process-related GO terms for biological regulation and regulation of transcription, while TSSL genes were enriched for plasma membrane-related cellular component GO terms. Based on functional annotations, there were 10 stem-preferred TFs from TALE, HD-ZIP, B3, YABBY, WRKY, AP2, bHLH protein families (***Table S3***). Also, there are 15 Andropogoneae family-specific stem-preferred genes (homologs are only found in *Sorghum bicolor, Saccharum spontaneum, Miscanthus sinesis*, and *Zea mays*), in which 10 of them are “orphan” genes without detectable homologues in other lineages for any previously annotated gene among eukaryotes, and are unique to a species and/or phylogenetic clade. Previous reports have shown that orphan genes generally have lower expression levels but higher tissue-specific expression compared to phylogenetically conserved genes (Levine *et al.*, 2006; Donoghue *et al.*, 2011; Wu, Irwin and Zhang, 2011). Several hypotheses have been proposed to explain the regulation of these younger genes that lead to tissue-specific and/or stage-specific expression, including transposon insertion upstream of the transcription start-site, shared regulatory elements with older genes (possibly by gene overlap), association with a bidirectional promoter, or by being located within an intron (Maksakova and Mager, 2005; Toll-Riera *et al.*, 2008; Arendsee, Li and Wurtele, 2014; Sundaram and Wysocka, 2020). We explored the cis-regulatory elements in the putative promoter regions of these genes below.

**Figure 3:**
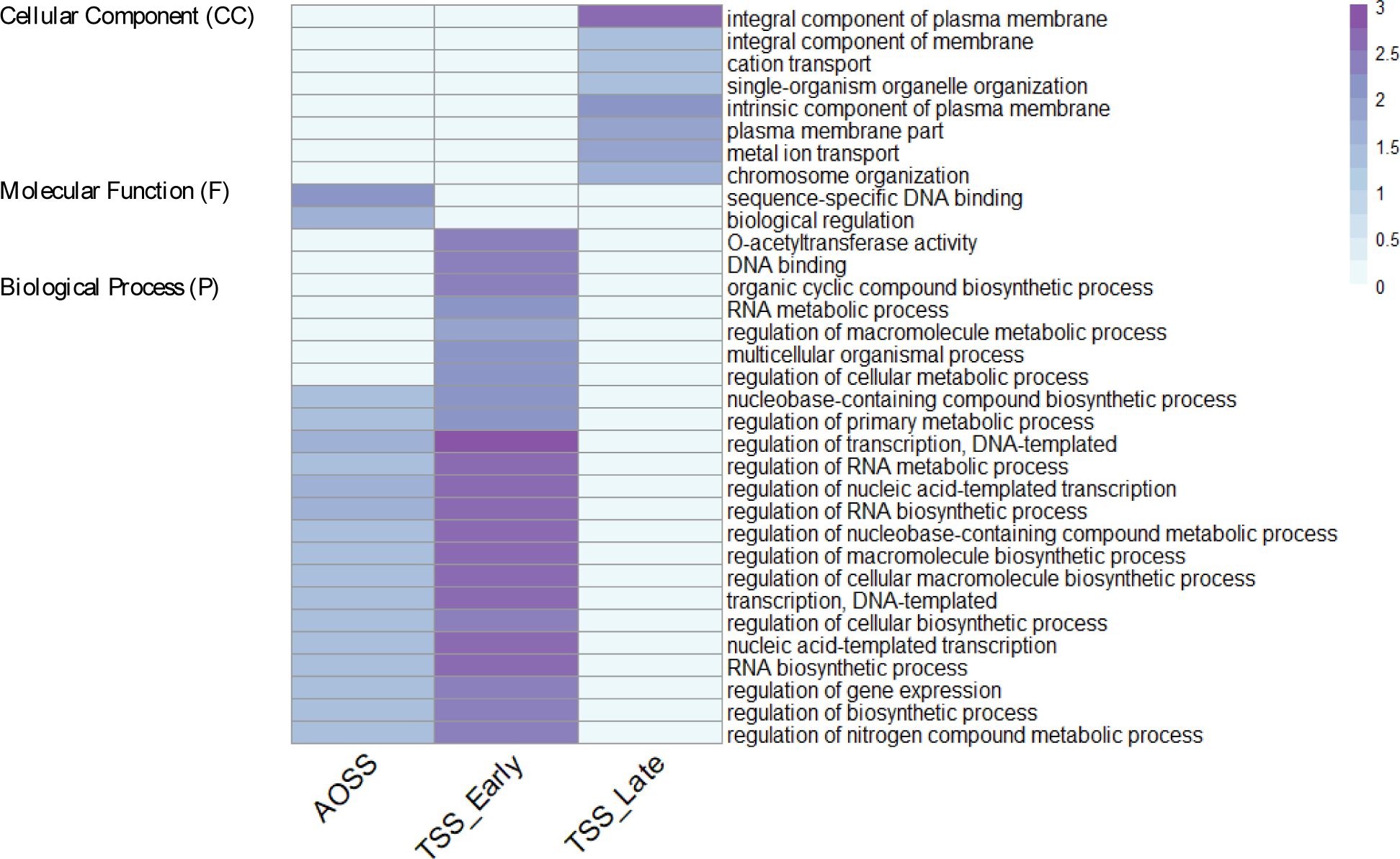
A set of GO terms enriched in AOSS, TSS-Early, and TSS-Late stem-preferred genes. A) Biological Processed GO Terms. B) Molecular Function and Cellular Component GO Terms. Significantly enriched GO terms are selected based on p-value < 0.05.

### Cis-regulatory element enrichment of AOSS, TSS-Early, TSS-Late stem-preferred genes

The three groups of stem genes (AOSS, TSSE, TSSL) were further explored in an attempt to uncover common and unique CREs that regulate these groups. Many CREs were common among promoters of genes of interest (GOI), including the orphan genes, across the three categories of stem-expressed genes (AOSS, TSS-Early, TSS-Late) (***Figure 4***), indicating that these genes are co-regulated by TFs that bind to the enriched CREs. These common CREs included binding sites for TF known to play roles in stem-specific functions such as bZIP, AT-Hook, HD-ZIP, GATA, MYB, SBP, TCP, WOX, which are previously reported to be present in vascular-specific genes (***Table S1***) (YH and SR, 2003; Ruiz-Medrano *et al.*, 2011; Le Hir and Bellini, 2013; Zhang *et al.*, 2014). Interestingly, the AP2 CRE was uniquely enriched in the AOSS group of genes, while the CPP CRE was strongly enriched in the TSSL group of genes compared to the AOSS and TSSE groups (***Figure 4***). This result suggests that TFs binding to the AP2 and CPP CREs might have an important role in the specific regulation of genes in the AOSS and TSSL groups.

**Figure 4:**
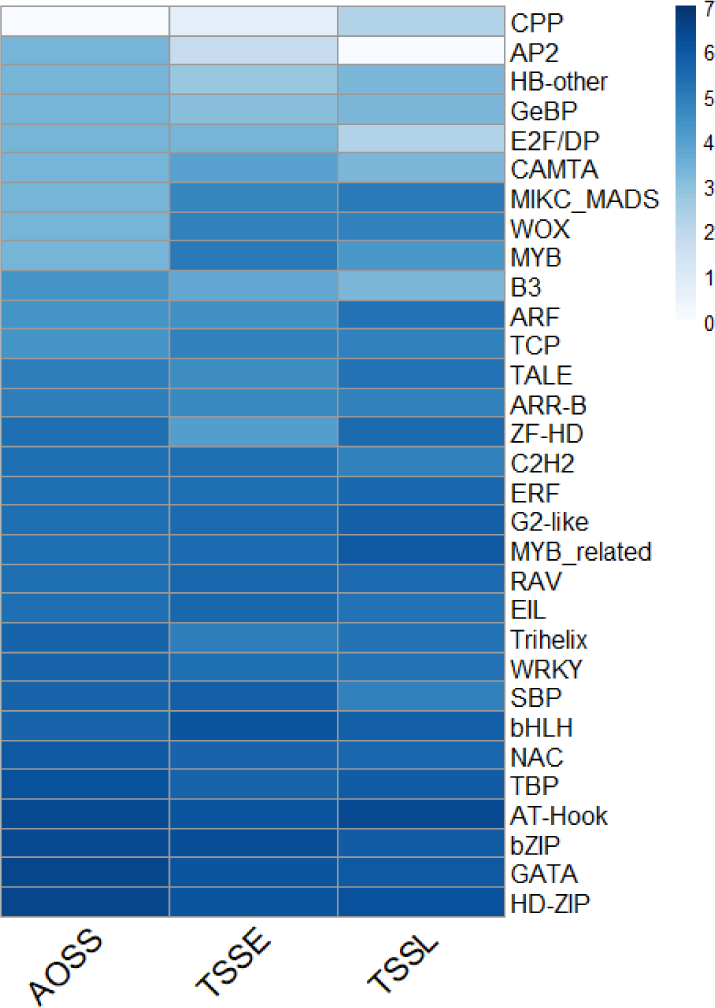
A set of unique cis-element motifs enriched in AOSS, TSS-Early, and TSS-Late gene promoters. The enrichment of cis-element motifs is identified based on over-representation of CREs in the 1.5 kb upstream promoters of stem-preferred genes, comparing to the whole genome (fold change > 1.1). The color intensity in the heatmap refers to percentage of stem-preferred sequences with enriched CRE.

### Gene regulatory networks of AOSS, TSS-Early, TSS-Late stem-preferred genes

Differential gene expression over development is controlled by temporal changes in TF binding to promoter elements. It is not only the timing of a TF’s expression, but also its temporal promoter occupancy that determines time evolved gene expression (Para *et al.*, 2014). However, extensive patterns of TF occupancy in developmental contexts are not known for sorghum (i.e. ChiP-Seq or DAP-seq). Therefore, elucidating the dynamic properties of cis-regulatory networks from available spatial-temporal datasets is useful in understanding how dynamic expression states arise. To explore which TFs regulate the expression of stem-specific genes, and examine whether stem-expressed genes are differentially regulated over time, gene regulatory networks (GRNs) were constructed for each group of stem-expressed genes at each developmental stage. GRNs were constructed by layering TF binding predictions onto gene co-expression networks. We generated 15-temporal stem-preferred gene regulatory networks (5 temporal GRNs for AOSS, TSS-Early, TSS-Late). See ***Data S1*** for list of stem-preferred genes and TFs and unique directed TF–target gene interactions for each of 15 GRNs.

GRN analysis of AOSS, TSS-Early, TSS-Late genes along with their co-expressed TFs revealed that the connectivity of each network has a positive correlation with the corresponding out-node degree, and that the out-node degree ‘*x’* in each of the networks follows a power-law distribution y=ax^b with an average exponent *b* of 0.182±0.064, 0.169±0.064, 0.23±0.07 for AOSS, TSS-Early, TSS-Late, respectively (***Table 3***). The connectivity of each network has a positive correlation with the corresponding out-node degree (AOSS: 0.661±0.205, TSSE: 0.653±0.127, TSSL: 0.629±0.229). This suggests that the GRNs have the properties of robust biological networks, which tend to contain scale-free properties such as having few nodes with a very large number of regulatory edges, or “hubs.” These hub genes are absent in random networks, for which the degree of all nodes is close to the average degree according to network theory (Cohen, Havlin and ben-Avraham, 2002; Barabási and Oltvai, 2004). Thus, the hubs in our GRNs potentially co-regulate many of the stem-expressed genes (***Figure 5***). In the stem GRNs, each TF is associated with an average of 4 genes during the early stages and 15 genes during the late stages (***Table 3***). Similarly, multiple TFs can regulate the expression of a single gene, and in the stem GRNs, each target gene is associated with an average of five regulatory TFs during the early stages compared to 18 regulatory TFs in the late stages. These results demonstrate that stem-expressed genes become increasingly co-regulated over development.

**Table 3:** Network characteristics in the Sorghum BTx623 spatial-temporal stem-preferred gene regulatory network. To characterize the topology of the stem-preferred gene’s temporal GRNs, each network’s topological characteristics were computed by NetworkAnalyzer in Cytoscape.

| **AOSS** |  |  |  |  |  |
| --- | --- | --- | --- | --- | --- |
|  | Juvenile | Vegetative | Floral Initiation | Anthesis | Grain Maturity |
| Total No. of genes in the network<br>(stem genes + TF) | 7+751 | 7+731 | 8+871 | 751+8 | 8+724 |
| Total No. of regulatory edges |  |  |  |  |  |
| Median no. of connections per gene |  |  |  |  |  |
| In-Degree | 6 | 4 | 28 | 14 | 12 |
| Out-Degree | 5 | 4 | 21 | 15 | 11 |
| Shortest path length | 13.4 | 16.6 | 3.7 | 4.5 | 4.7 |
| Clustering coefficient | 0.306 | 0.292 | 0.215 | 0.234 | 0.225 |
| Connectivity of out-node distribution |  |  |  |  |  |
| Degree exponent | 0.182 | 0.082 | 0.248 | 0.211 | 0.217 |
| R=Squared | 0.661 | 0.203 | 0.679 | 0.653 | 0.653 |
| **TSS-Early** |  |  |  |  |  |
| Total No. of genes in the network<br>(stem genes + TF) | 42+779 | 38+740 | 38+875 | 15+751 | 13+724 |
| Total No. of regulatory edges |  |  |  |  |  |
| Median no. of connections per gene |  |  |  |  |  |
| In-Degree | 6 | 4 | 29 | 14 | 12 |
| Out-Degree | 5 | 4 | 20 | 15 | 11 |
| Shortest path length | 13.4 | 16.8 | 3.7 | 4.5 | 4.7 |
| Clustering coefficient | 0.305 | 0.285 | 0.211 | 0.233 | 0.223 |
| Connectivity of out-node distribution |  |  |  |  |  |
| Degree exponent | 0.169 | 0.092 | 0.252 | 0.204 | 0.237 |
| R=Squared | 0.653 | 0.392 | 0.675 | 0.589 | 0.711 |
| **TSS-Late** |  |  |  |  |  |
| Total No. of genes in the network<br>(stem genes + TF) | 2+460 | 3+729 | 4+870 | 4+750 | 5+723 |
| Median no. of connections per gene |  |  |  |  |  |
| In-Degree | 6 | 4 | 29 | 14 | 12 |
| Out-Degree | 5 | 4 | 21 | 14 | 11 |
| Clustering coefficient | 0.290 | 0.291 | 0.214 | 0.233 | 0.225 |
| Connectivity of out-node distribution |  |  |  |  |  |
| Degree exponent | 0.230 | 0.071 | 0.245 | 0.215 | 0.221 |
| R=Squared | 0.629 | 0.150 | 0.674 | 0.649 | 0.686 |

**Figure 5:**
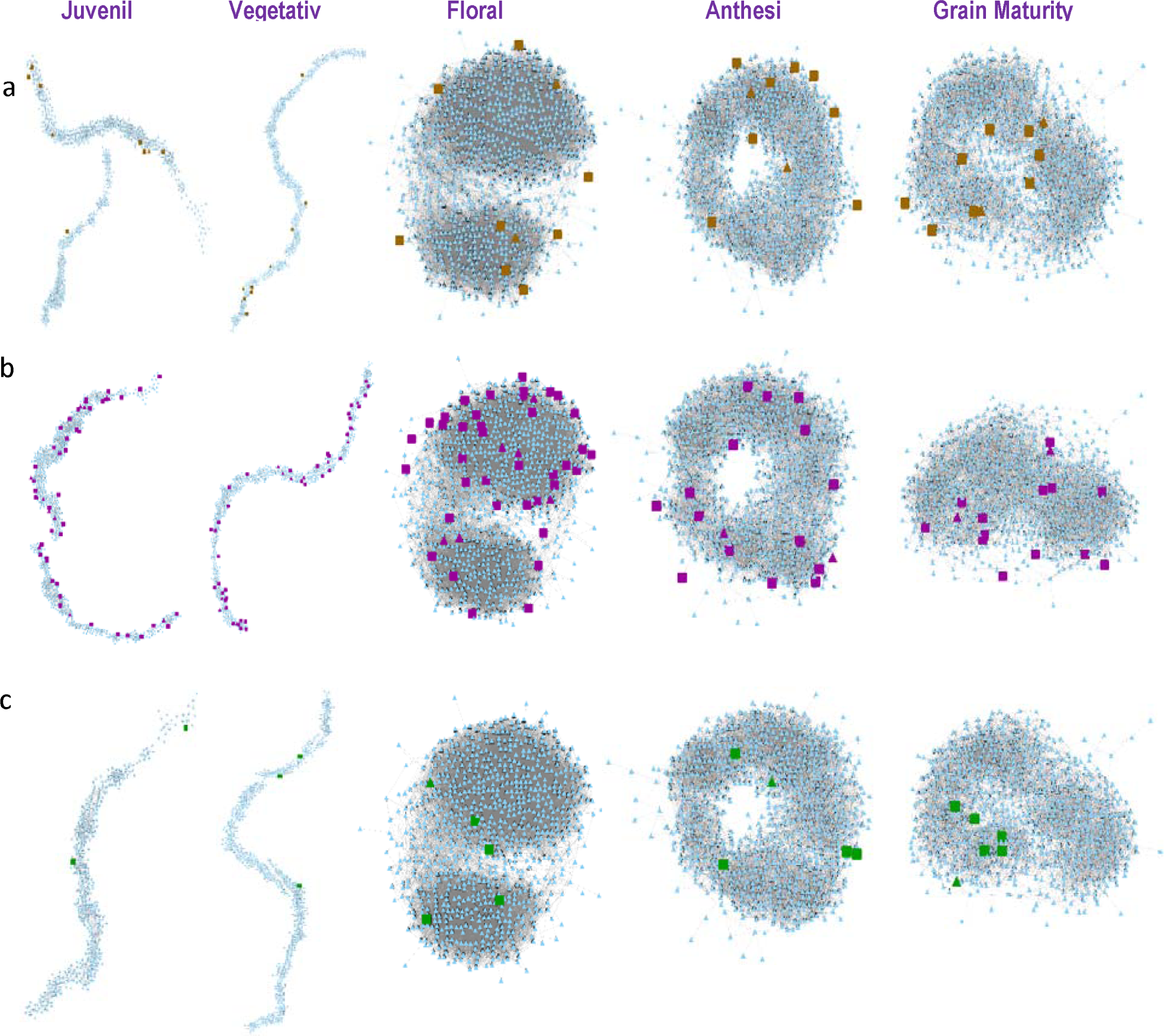
Temporal gene regulatory network of a) AOSS, b) TSSE, c) TSSL genes through five developmental stages in BTx623 sorghum. Edges with Pearson Correlation >= 0.7 and MR >= 0.7 is shown here.

#### TF Hubs enriched for known stem genes

We evaluated the hubbiness of the TFs in our GRNs to identify TFs that exert a large amount of control over the entire network. Hubs are defined here as highly connected TF genes (top 25% h-index) (See ***Table S4*** for list of TF hubs in each network). The most frequent TF families present in the networks are bHLH, bZIP, ERF, C2H2, G2-like, GATA, HD-ZIP, NAC, MYB, MYB-related, TALE, Trihelix, and WRKY. TFs previously reported as important regulatory genes in stem tissue are present in the networks, including known stem TF hubs such as PHB, ATHB15, APL, VND7, KNAT1, MYB42, and MYB52 (McConnell *et al.*, 2001; Ye, 2002; Emery *et al.*, 2003; Kubo *et al.*, 2005; Floyd, Zalewski and Bowman, 2006; Hoon Jung and Mo Park, 2007; Zhong *et al.*, 2008; Zhou, Sebastian and Lee, 2011; De Rybel *et al.*, 2016; Hennet *et al.*, 2020). Radial patterning of the vascular bundles is established by TF hubs from class III HD-ZIPs, PHB, ATHB15 (McConnell *et al.*, 2001; Emery *et al.*, 2003; Floyd, Zalewski and Bowman, 2006). NACs are reported to be TF hubs that can induce xylem cell differentiation (Kubo *et al.*, 2005; Zhong *et al.*, 2008). For example, VND7 (NAC TF Family) initiates metaxylem vessel differentiation and induces hierarchical gene expression network by inducing other TFs such as MYBs, which in turn function as secondary and tertiary regulators of genes related to secondary cell wall thickening during xylem cell differentiation (Ye, 2002; Hoon Jung and Mo Park, 2007; Zhou, Sebastian and Lee, 2011; De Rybel *et al.*, 2016).

Next, we compared hubbiness of TFs over development to identify significant TF genes that change their hub property status relevant to early and late developmental phases. Our expectation was there might be different set of critical TFs that drive stem-specific gene expression during early and late development. There were 22 and 22 TFs that acted as early TF hubs (act as a TF hub only during early stages) for AOSS and TSSE genes, respectively. Likewise, there were 19 and 21 late TF hubs (act as a TF hub only during late stages) for AOSS and TSSL genes, respectively (***Table S5***). We examined whether any of these early and late TF hubs are unique to AOSS vs. TSSE/TSSL genes. However, the results showed all early and late TF hubs are critical TF hubs for AOSS, TSSE, TSSL genes during early and late stages, respectively. Likewise, genes that remain a functionally conserved TF hub in all GRNs during early and late phases also tend to be TF hubs for all AOSS, TSSE, TSSL GRNs. This could mean stem-specific and transiently stem-specific genes are likely co-regulated by the same higher-level transcriptional program during respective early and late developmental phases.

The stem-preferred spatial-temporal GRNs exhibit different transcriptional regulatory programs that activate the stem-preferred genes over time (***Table 3, Figure 5***), revealing active TFs that temporally drive stem-specific expression during early and late development. To further explore this possibility, we compared early (juvenile, vegetative) regulators to late (floral, anthesis, grain) regulators of stem-preferred genes to identify stage-specific TFs. There were six TFs predicted to regulate at least one AOSS or TSSE gene during the early stages (SP-Early regulators: AIB/Sobic.003G272200 (regulatory edges to 1 AOSS and 3 TSSE), Sobic.004G283300/Integrase-type DNA binding superfamily protein (regulatory edges to 3 TSSE), PDF2/ Sobic.007G065300 (regulatory edges to 3 AOSS and 2 TSSE), bHLH/Sobic.007G189400 (regulatory edges to 1 AOSS and 4 TSSE), MYB61/Sobic.009G036500 (regulatory edges to 2 TSSE), and WRKY33/Sobic.009G100500 (regulatory edges to 2 TSSE)). Only one TF was identified that uniquely regulates the expression of stem-specific genes during the late developmental stages in the AOSS and TSSL gene groups (SP-Late regulator: RSM2/ Sobic.009G235500 (regulatory edges to 6 AOSS and 4 TSSL)).

Next, we tested whether there are unique TFs belonging to different CREs that turn on AOSS genes vs. TSSE or TSSL genes. Therefore, we tested whether there are: a) unique TF regulators specific to AOSS genes during early stages; b) unique TF regulators specific to AOSS genes during late stages; c) unique TF regulators specific to TSSE genes during early stages; and d) unique TF regulators specific to TSSL during late stages. There were no TFs unique to the AOSS and TSSL networks over time. However, three TFs, MYB61, WRKY33, and Sobic.004G283300, are predicted to uniquely regulate TSSE genes during juvenile and vegetative networks.

### Comparison of gene regulatory networks of stem-preferred genes between BTx623 and Della cultivars

To examine if there is conserved regulation of stem-specific gene expression among grain and sweet sorghum varieties we compared the temporal stem-specific GRNs between *cv.* BTx623 (grain) and *cv.* Della (sweet) using transcriptome profiles from McKinley et al. (2018) and McCormick et al. (2018). Comparative analysis revealed little consensus among regulatory edges (average consensus of 1.8% for AOSS, 1.8% for TSSE, 1.3% for TSSL) between BTx623 and Della GRNs at comparable developmental stages (***Figure 6, Figure 7, Figure 8***). Thus, GRN analysis suggests that there are different transcriptional regulatory programs regulating the expression of stem-specific genes in these two cultivars. While the majority of regulatory edges directly connected to the stem-specific genes are variable between the two cultivars, it is interesting to note that whole network analysis revealed that the same TF hubs are conserved, with an average consensus of 38.9%, 39.5%, and 46.0% in the AOSS, TSSE, and TSSL, respectively.

**Figure 6:**
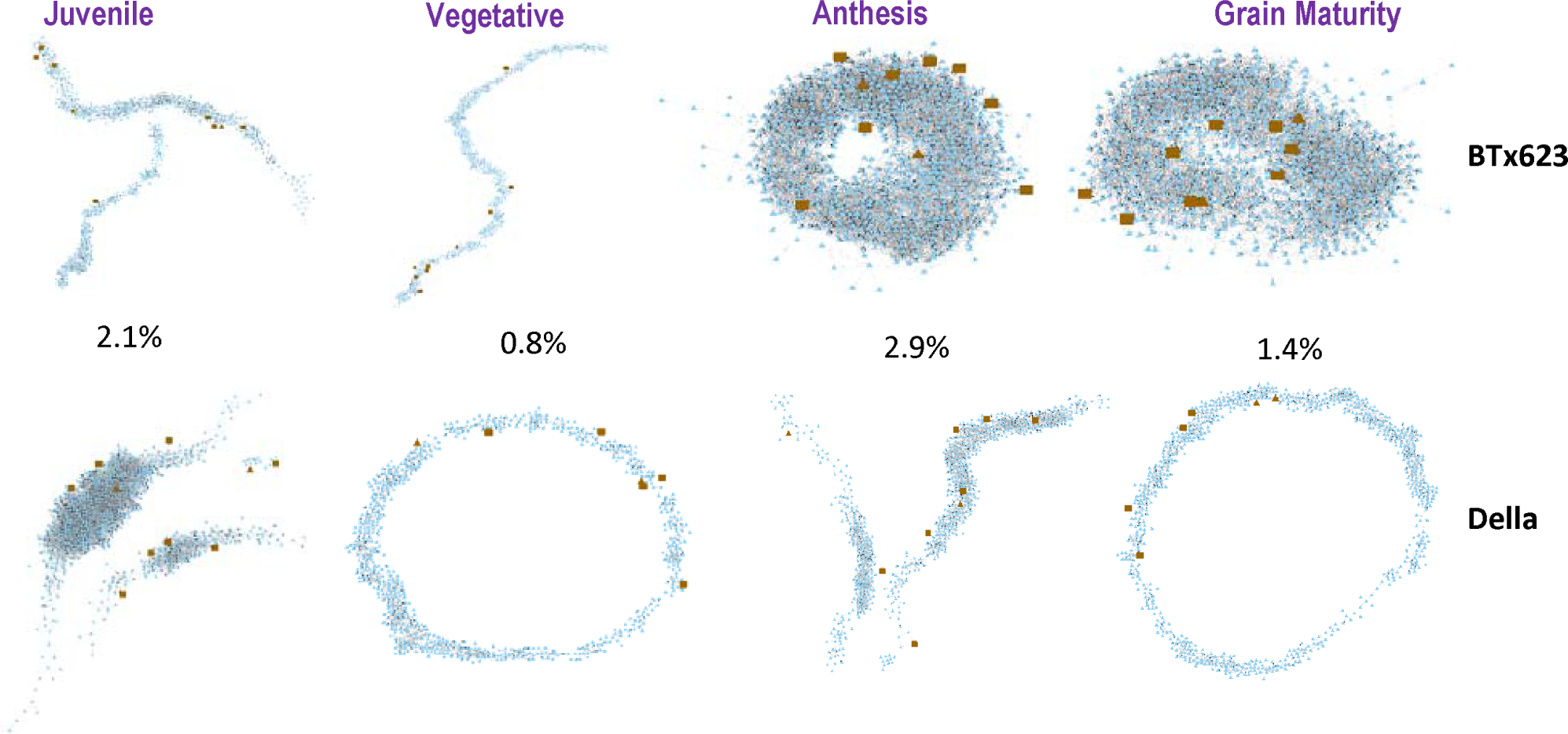
Temporal gene regulatory network of AOSS genes through development in BTx623 and Della cultivars. Edges with Pearson Correlation >= 0.7 and MR >= 0.7 is shown here. The percentage of shared regulatory edges are listed.

**Figure 7:**
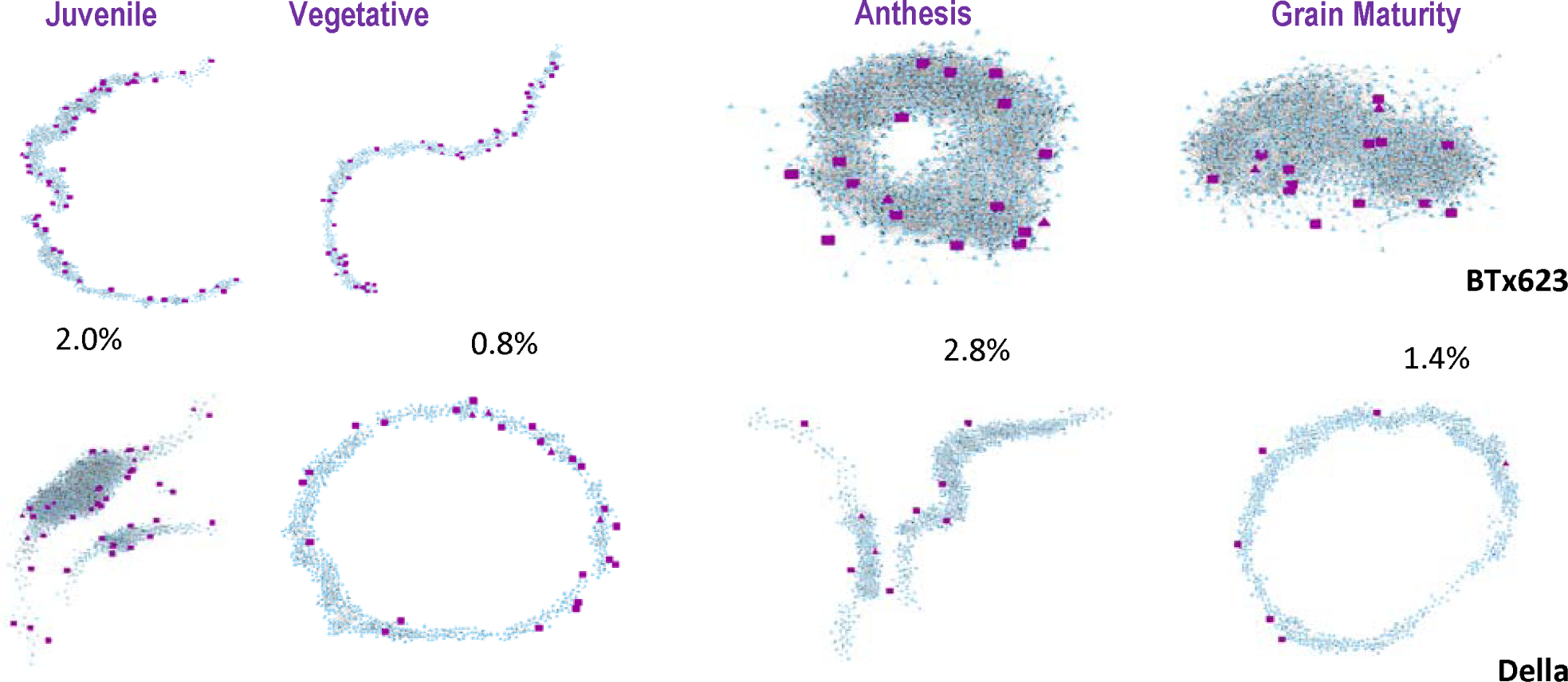
Temporal gene regulatory network of TSSE genes through development in BTx623 and Della cultivars. Edges with Pearson Correlation >= 0.7 and MR >= 0.7 is shown here. The percentage of shared regulatory edges are listed.

**Figure 8:**
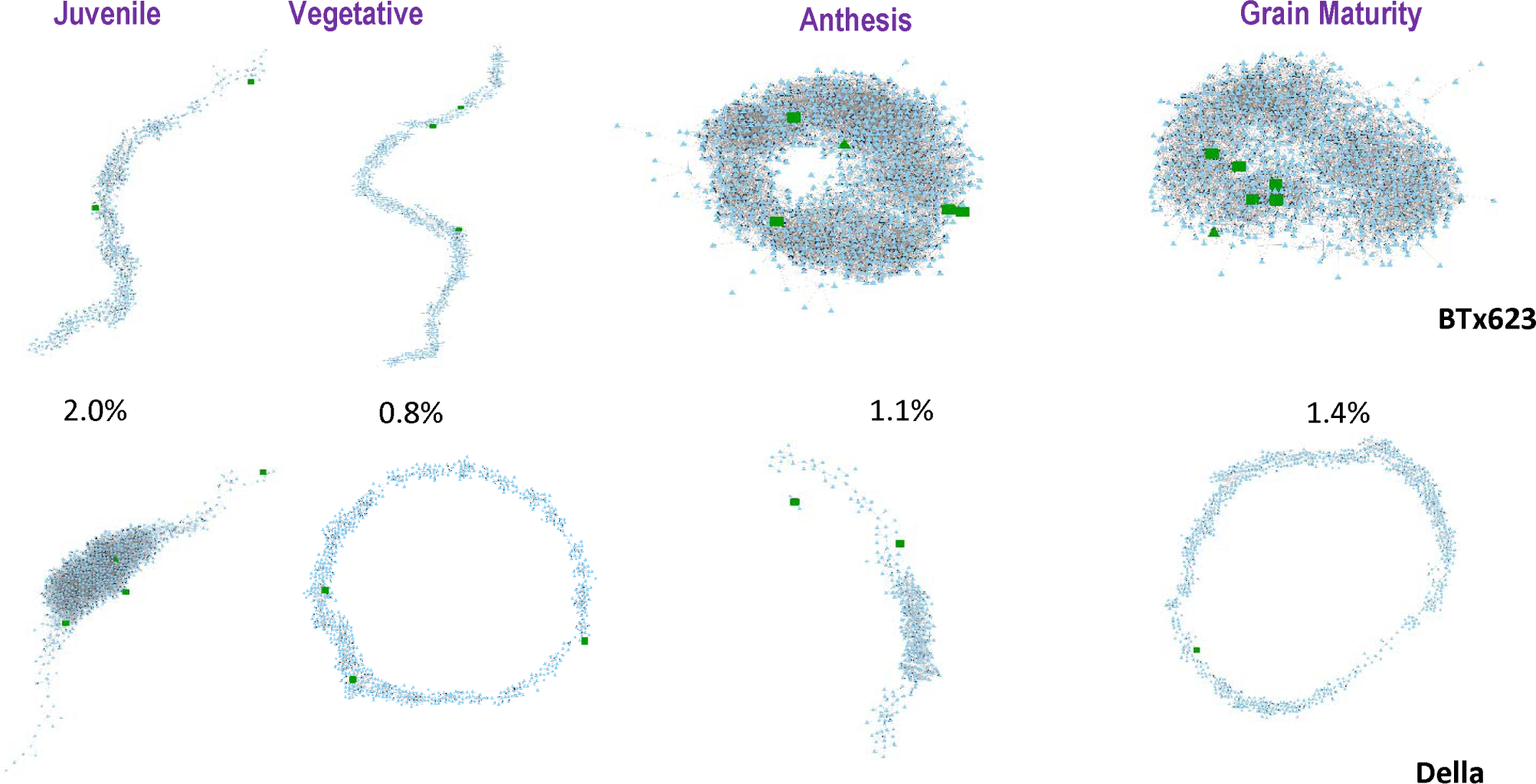
Temporal gene regulatory network of TSSL genes through development in BTx623 and Della cultivars. Edges with Pearson Correlation >= 0.7 and MR >= 0.7 is shown here. The percentage of shared regulatory edges are listed.

To further investigate the variation in regulatory edges in the stem GRNs between cultivars, we explored the genetic sequences of stem expressed GOIs. Various events can alter the expression of a gene, such as insertions, deletions, and point mutations in either the gene of interest or the transcription factors (TFs) that regulate that gene. Regulatory DNA variants adjacent to or within tissue-specific genes or their corresponding transcriptional regulators may influence gene expression patterns in different cultivars. While many mechanisms can regulate these genes differently (i.e. involvement of various cell- and species-specific non-coding cellular factors such as post-transcriptional regulators and post-translational modifications), it is reasonable to examine the contribution of cis-regulatory element variants for divergent regulation. This was elucidated by estimating genetic variation between BTx623 and Della *cvs*, for insertions, deletions, and point mutations. Several single nucleotide variations (SNVs) and/or structural variations (SVs) between BTx623 and Della were found in the gene body, 1.5 kbp upstream, and/or 500 bp downstream regions for a number of stem-expressed genes in the AOSS, TSSE, and TSSL gene groups. The list of SNVs and SVs found in each of these groups is in ***File S1*** and ***File S2***, respectively. For example, within the AOSS group of genes, six out of ten genes had at least one SNV and/or SVs in the gene body, 1.5 kbp upstream, or 500 bp downstream region between BTx623 and Della. Of these genes, four contained these mutations within a CRE (***Table 4*** and ***Figure S3***) in their putative promoter regions. For example, both Sorghum homologous of KNAT1 (Sobic.001G106000 and Sobic.001G106200) had multiple CREs with SNVs, including *NAC, TBP, SBP, AT-Hook, GATA, ARR-B, bZIP, C2H2, MADS-Box* (***Table 4***).

**Table 4:**
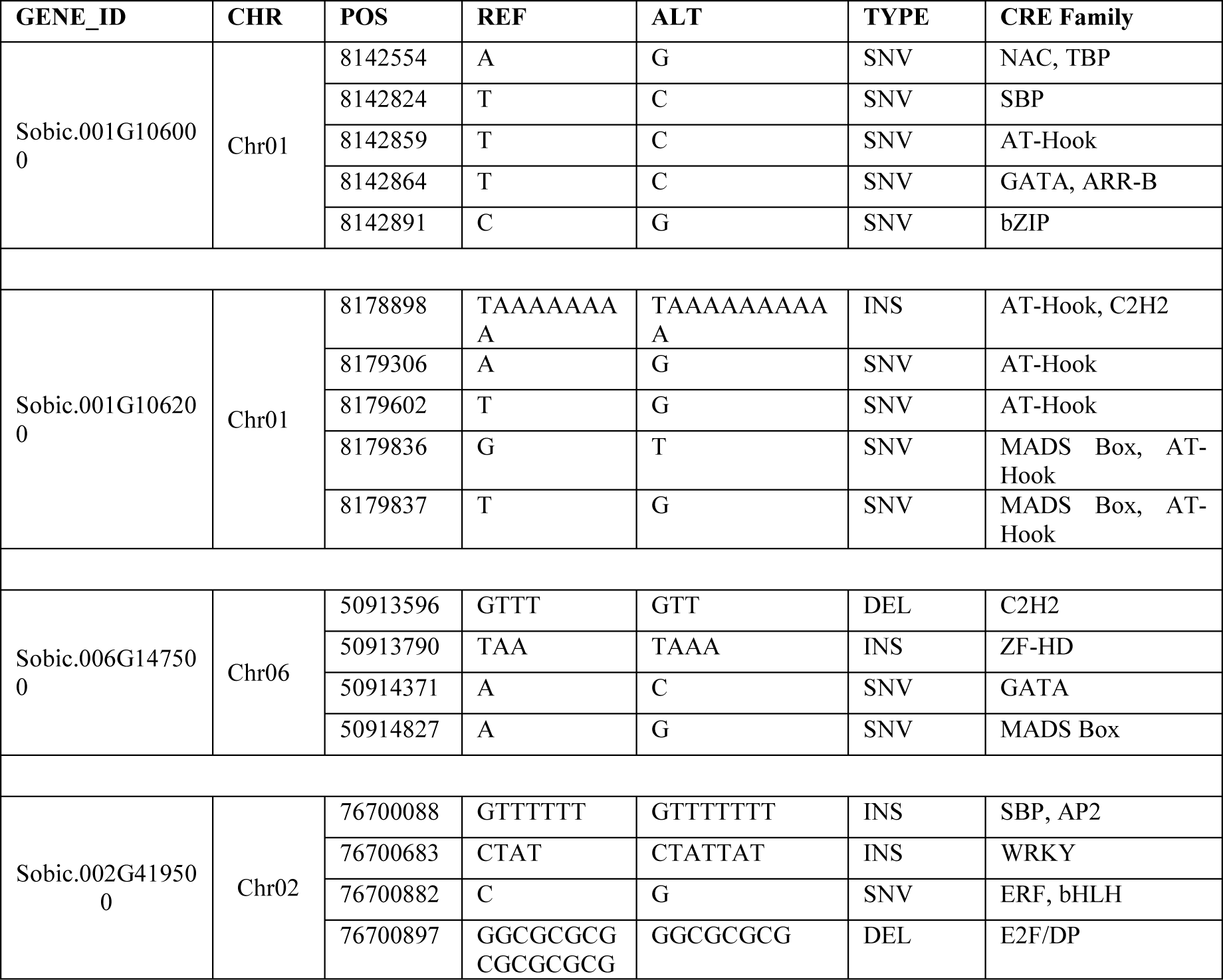
A table of SNVs and SVs within cis-regulatory elements in 1.5 Kbp upstream non-coding region of stem-specific genes with significant genetic alterations.

### Genetic variation in hierarchical regulators may contribute to the differential activation of stem-specific genes among Sorghum cultivars

In cases where stem-expressed GOIs have different GRNs but no genetic variations between the cultivars, we explored if genetic alterations exist in the genes encoding TFs predicted to regulate the stem-specific GOIs. Again, several single nucleotide variations (SNVs) and/or structural variations (SVs) between BTx623 and Della were found in the gene body, 1.5 kbp upstream, and/or 500 bp downstream regions for a number of TFs predicted to regulate stem-expressed genes in the AOSS, TSSE, and TSSL gene groups (***File S1*** and ***File S2***). For example, 271 out of 381 total TFs across the AOSS GRNs had at least one SNV and/or SV within the gene region under investigation. Of those, 167 TF genes (136 are considered network hubs) had genetic alterations inside a CRE within their putative promoter region. Key TF hubs with variations include PHB, ATHB15, KNAT1, GATA4, ABF4, AGL7, HB-7, TGA4, VIP1, MYB48, FMA, GBF3, and LHY1. TF hubs with genetic variations likely contribute to the observed global differences in stem GRNs between cultivars. In a specific example, Sobic.006G026300 is one of most stem-specific and highly expressed genes in Sorghum *cv.* BTx623. ***Figure 10*** shows TFs predicted to regulate Sobic.006G026300 over the five developmental stages. The network prediction for Sobic.006G026300 in *cv BTx623 and cv Della* showed different regulation patterns (***Figure 9***). The genomic region of this gene is highly conserved with no observed mutations within the gene body or up- or down-stream non-coding regulatory regions (***Figure S2***). One explanation could be genetic variations present in the genes encoding the TF regulators between cultivars. ***Table 5*** contains list of SNVs and SVs for some of the predicted TF regulators of Sobic.006G026300, including NAC1 (Sobic.008G164800), a regulator during the vegetative – grain stages (***Figure 10***), that has several significant structural variation, namely TBP (INS), bZIP (INS/RPL), AT-Hook (INS/RPL). Likewise, the TF bHLH93 (Sobic.001G513700), a regulator during the juvenile-vegetative stages (***Figure 10***), has structural variations within the RAV (DEL), MADS-Box (DEL), G2-like (DEL), ERF (INS), and C2H2 (INS) CREs, and the TF ABF4 (Sobic.002G225100), also a regulator during the juvenile stage, has a 24 bp and a 29 bp deletion next to each other flanking WRKY, MYB, MYB-related CREs, and 21 bp insertions within the bZIP and bHLH CREs in its putative promoter region. Because changes in stem-specific gene expression between species could also arise by diversification of master-regulators, altering hub expression can cause large-scale network changes, potentially driving the different tissue-specific gene regulatory networks we observed.

**Table 5:**
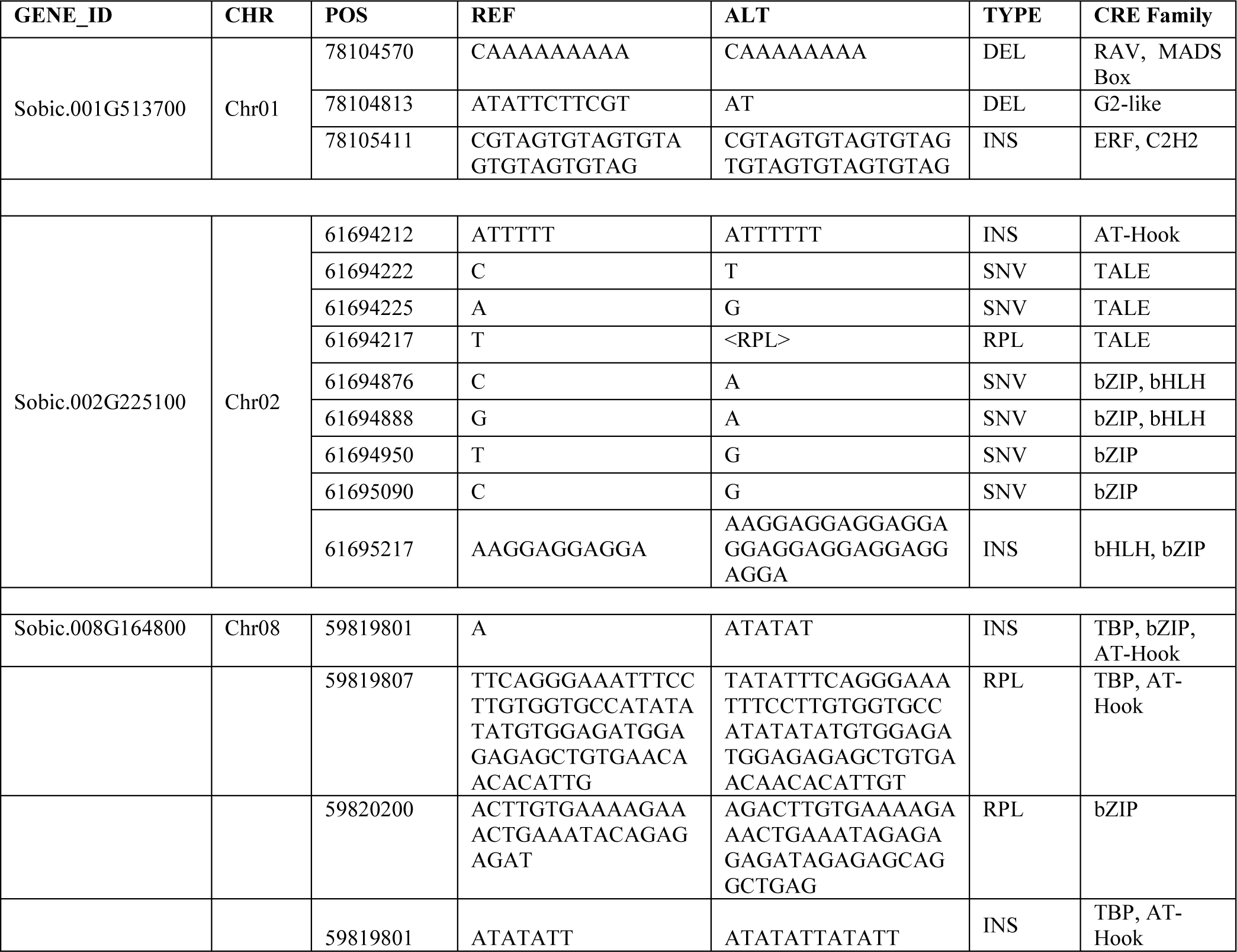
Genetic alterations within cis-regulatory elements in 1.5 Kbp upstream non-coding region between Della and BTx623 cultivars for predicted regulatory TFs of Sobic.006G024800 during a) Early stage and b) Late stage.

**Figure 9:**
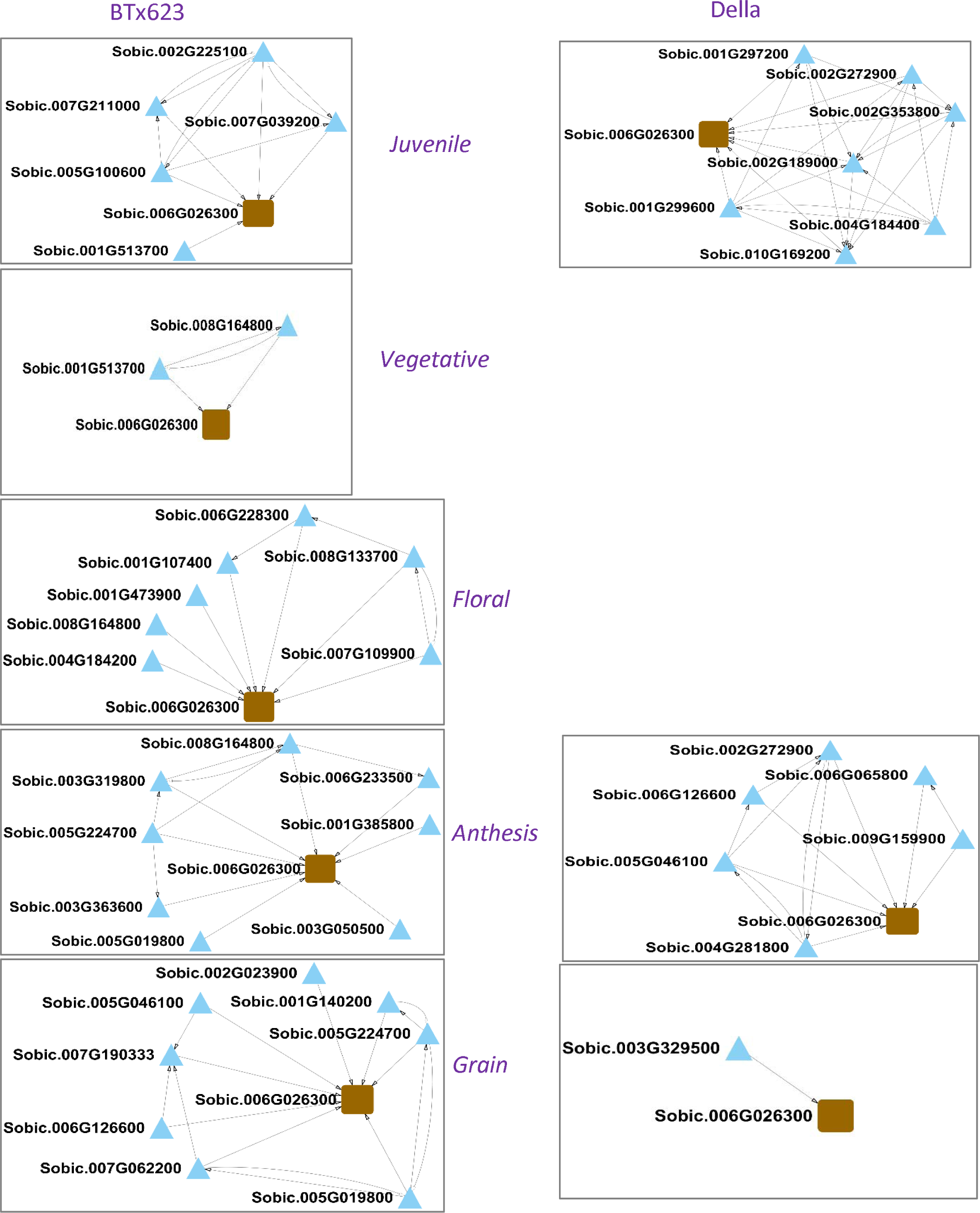
Predicted gene regulatory networks of Sobic.006G26300 (one of most stem-specific and highly expressed gene) for BTx623 and Della for comparable time points.

**Figure 10:**
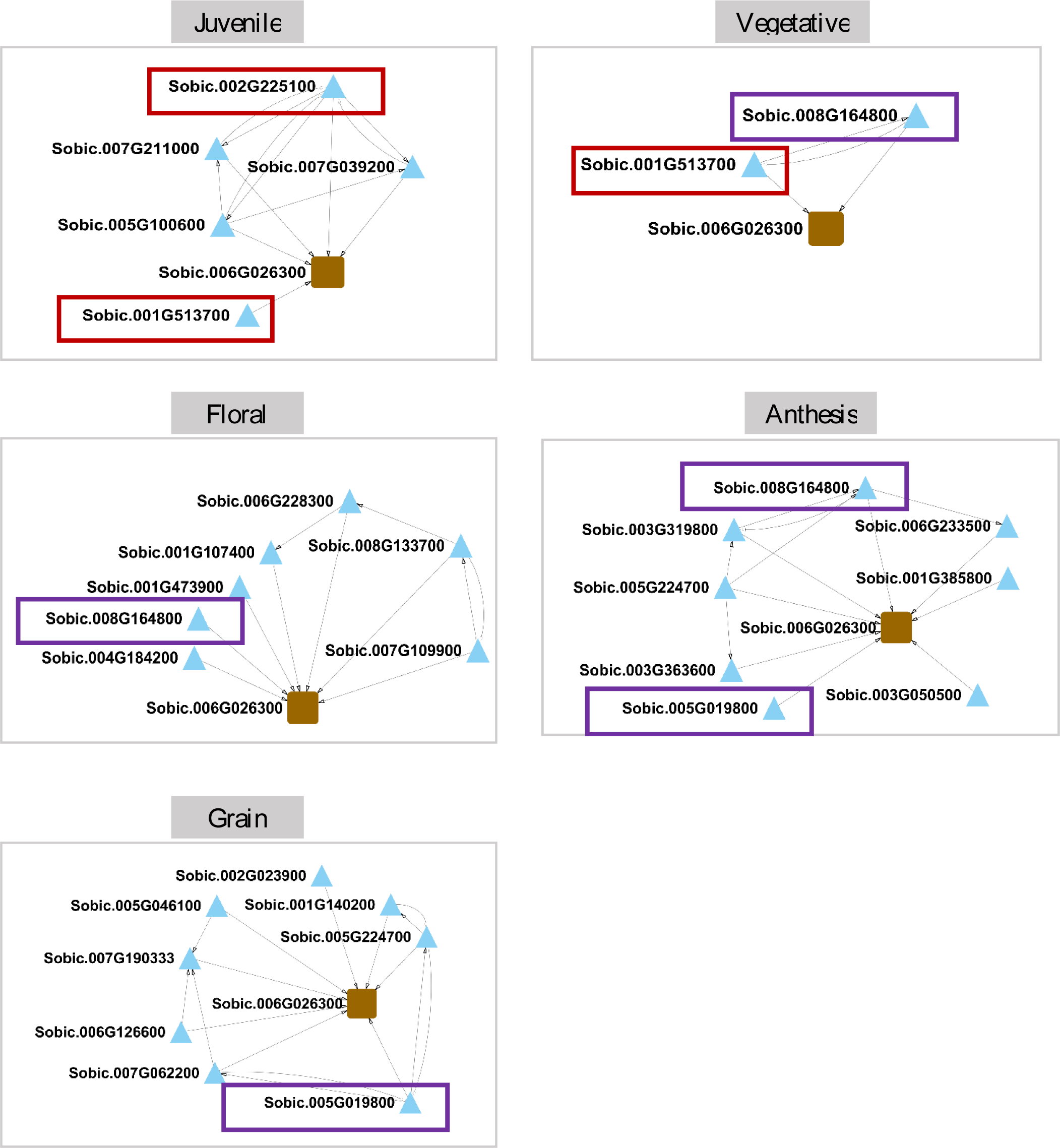
Predicted regulatory TFs of Sobic.006G024800 during a) Early stage and b) Late stage. The regulatory interactions shown here are predicted at PCC & MR >= 0.9.

## CONCLUSIONS

Using *in-silico* analysis of spatial-temporal transcriptome datasets, we report a set of “stem-specific” genes that are always expressed in the stem throughout development, and “stem-preferred” genes that are transiently stem-specific over development. This includes genes known to be involved in vascular tissue development and/or function such as KNAT1, HB7, MYC2, CLV3, SUVR1, WRKY12, AGO1, PLC2 (Hoon Jung and Mo Park, 2007; Martínez-Navarro *et al.*, 2013; De Rybel *et al.*, 2016; Hennet *et al.*, 2020). The identification of common cis-regulatory elements within the promoters of stem-specific and stem-preferred genes suggest co-regulation, which was further explored using predictive GRNs. Network analysis revealed that GRN topology changes over time, indicating that different transcriptional regulatory programs are responsible for the temporal activation of the stem-expressed genes. However, TF hubs appear to be conserved across the temporal stem GRNs and include known regulators of stem genes as well as TFs not previously implicated in stem development/function that merit further investigation.

The analysis of temporal stem GRNs between grain and sweet sorghum varieties revealed striking differences in the regulatory landscape, which is not entirely surprising given the considerable phenotypic differences between grain and sweet sorghums. The sorghum BTx623 cultivar is an early maturing grain sorghum genotype with short stature, which is phenotypically distinct from the tall, late-maturing Della cultivar (sweet sorghum genotype), typically grown for stem sugars or high biomass yield (Jiang *et al.*, 2013; Cooper *et al.*, 2019). BTx623 encodes recessive alleles of the dwarfing gene *Dw1* (Hilley *et al.*, 2016) that encodes a protein involved in brassinosteroid signaling (Hirano *et al.*, 2017) and *Dw3*, a gene that encodes an ABCB1 auxin efflux transporter (Multani *et al.*, 2003; Knöller *et al.*, 2010), whereas Della is dominant for both dwarfing loci. The difference in dwarfing genotype results in differences in internode lengths and the ratio of node/internode tissues in BTx623 and Della. Grain sorghums develop treachery elements during xylem formation earlier than sweet sorghums, and sweet sorghums accumulate more stem sugars compared to grain sorghum (Jiang *et al.*, 2013; Cooper *et al.*, 2019). Grain sorghums usually contain less carbohydrate in the upper internodes at grain maturity compared with sweet sorghum varieties, due to mobilization of carbon from stems during grain filling. The divergence of stem related genes between the two types has previously been suggested from genetic variation analysis (Jiang *et al.*, 2013; Cooper *et al.*, 2019), and the present study suggests genomic divergence in stem-expressed genes based on their developmental transcriptomes and regulatory programs.

Genomic perturbations such as single nucleotide variants and large structural variants can lead to an altered function or expression of TFs involved in stem-specific functions that lead to downstream changes in TF activity (e.g., HD-ZIP, MYB, NAC, TALE). Significant nucleotide sequence variation in a canonical TF binding motif in a promoter or enhancer region affects the prediction of TF-target edges in a regulatory network. Such perturbations in CRE sites within putative promoter regions of stem-expressed genes resulted in different GRNs for *BTx623* and *Della* cultivars. However, further studies are needed to confirm “real” TF-target interactions through experiments such as yeast-one hybrid (Y1H) (Arda and Walhout, 2009; Fuxman Bass, Reece-Hoyes and Walhout, 2016) or DAP-seq (O’Malley *et al.*, 2016). Nonetheless, the current study provides a starting point for more in-depth experiments to assess the extent to which these promoter elements are tissue-specific through targeted mutagenesis, for example, using tissue-specific genome modification (Schürholz *et al.*, 2018; Decaestecker *et al.*, 2019; Shi *et al.*, 2019; Ali, Mahfouz and Mansoor, 2020). This will greatly benefit efforts to fine-tune manipulation of tissue-specific and developmental-stage specific gene expression of transgenes required to genetically engineer sorghum stems to accumulate oil and elevate its status as a biofuel crop.

## Methods

### RNA-Seq data set representing different tissues over 5 development time points

We used a publicly available *Sorghum bicolor* transcriptomic dataset containing expression profiles of 38 tissue samples (McCormick *et al.*, 2018b) representing four plant organs (stem, leaf, root, and reproductive structures) over five developmental stages (juvenile, vegetative, floral initiation, anthesis, and grain maturity). This dataset was used to obtain a tissue-specific expression atlas (***Table 1***).

The quality cleaned raw reads of the 38 RNA-seq samples, *Sorghum bicolor (cv BTx623)* reference genome (S. bicolor v3.1.1), and genome annotations were downloaded from Phytozome (phytozome.jgi.doe.gov). The reads were then mapped to the reference genome using the STAR/2.6.0c-aligner (Dobin *et al.*, 2013). Read numbers that mapped to each gene were counted by featureCounts v.1.4.5-pl under the subread/1.5.2 package (Liao, Smyth and Shi, 2014). The normalized gene expression levels were estimated as in transcripts-per-million (TPM) using modules from edgeR package (Robinson, McCarthy and Smyth, 2010) in R (R Development Core Team, 2019). In a given tissue sample at a given developmental stage, genes having more than five counts per million (CPM) were considered as expressed. To visualize close grouping of spatial-temporal samples, hierarchical clustering and principal component analysis (PCA) of the log-transformed and quantile normalized TPM counts was performed and visualized using ‘FactoMineR’ R package (***Figure 1***). This revealed that roots, stems, and leaves formed distinct clusters based on gene expression, while stem samples formed a cluster with the peduncle and panicle (***Figure 1*).** Grass peduncles are the specialized stem internodes formed after floral initiation that support the panicle, consistent with co-clustering with other stem tissues. Peduncle tissue and the first panicle tissue sample was collected from BTx623 approximately 7 days after floral initiation when the peduncle/panicle was in an early stage of development. During early stages of inflorescence development, the rachis, the stem tissue of an inflorescence, constitutes a large portion of harvested panicle tissue. This helps explain why stem, peduncle and panicle tissues form a cluster.

### Identification of stem-preferred genes

We computed a tissue-specificity index to explicitly identify stem-specific genes expressed constitutively across all development stages or only during a specific developmental stage using tissue-specific RNA-seq gene expression profiles from McCormick et al. 2018. In this study, Tau Index (τ) (Kryuchkova-Mostacci and Robinson-Rechavi, 2016) was used to identify tissue-specific and broadly expressed genes. The index τ is defined as:

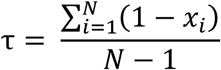

Where, *N* is the number of tissue types, and *x*_*i*_ is the expression profile component for a given gene in a given tissue, normalized by the maximal expression value of the given gene from all tissue samples considered. A custom R script adapted from (Kryuchkova-Mostacci and Robinson-Rechavi, 2016) was used to perform Tau index analysis (https://github.com/Madara-Dilhani/Tissue_Specificity). The τ values can vary from 0-1; genes with a τ value of 1 being absolute tissue-specific genes that are only expressed in a single tissue, and genes with a τ value of 0 are expressed in all tissues. Generally, tissue-specificity measures produce a bimodal distribution enabling identification of these groups, a) 0-0.2 tissue specificity (non/low specificity, housekeeping genes), b) 0.2-0.8 tissue specificity (intermediate specificity), c) 0.8-1.0 tissue specificity (high/absolute specificity). For example, “Housekeeping Genes” are often expressed in many (or all) tissues with no enriched expression for a particular tissue, thus the reported Sorghum housekeeping genes PP2A (Sobic.004G092500, Sobic.010G174800, Sobic.001G492300, Sobic.001G049400, Sobic.001G251700) and EIF4-α (Sobic.004G039400, Sobic.004G356900, Sobic.006G121100, Sobic.010G251100, Sobic.001G224400, Sobic.002G335600, Sobic.003G442400) had τ values < 0.2 (Sudhakar Reddy *et al.*, 2016; Nguyen, Eamens and Grof, 2018). A few previously reported maize stem-preferred gene homologs in sorghum were identified as highly stem specific (τ = 1); Glycosyl Hydrolase (Sobic.002G273500), AP2 (Sobic.003G203800, Sobic.008G149400), KNAT (Sobic.001G106000, Sobic.001G106200, Sobic.001G075101), RNI (Sobic.001G075101, Sobic.006G022500, Sobic.006G088820), WRKY (Sobic.004G298400, Sobic.006G166300), SUV (Sobic.004G218000), CLAVATA (Sobic.004G207000), BRX (Sobic.006G203200), CDC2 (Sobic.006G202100) (Hoopes *et al.*, 2019).

The mean gene expression (quantile normalized log-TPM) across replicates within an organ at a single developmental stage was determined. The τ index was calculated per each expressed gene at each developmental stage. The τ index is associated with its respective tissue with maximum expression. In this analysis, a gene was considered tissue-specific if it had a τ = 1. Following τ analysis, three groups were identified: 1) genes that are always tissue-specific throughout development, now called Always On Stem Specific (AOSS), 2) genes that are stem-specific during early developmental stages (juvenile and vegetative), now called Temporally Stem Specific Early (TSS-Early), and 3) genes that are tissue-specific during late developmental stages (floral initiation – grain maturity), now called Temporally Stem Specific Late (TSS-Late). The log_2_-TPM values of 3 categories of stem-specific genes were visualized in heatmaps generated using ggplot2 R package (***Figure 2***).

### Enrichment of biological functions and cis-regulatory elements

We identified sets of enriched gene ontology terms (GO-terms), and cis-regulatory elements (CREs) for each stem-specific gene set separately (AOSS, TSS-Early, TSS-Late). The GO enrichment tool (p-value p ≤ 0.01) at PlantTFDB v5.0 (http://planttfdb.gao-lab.org) was used to identify enriched GO terms (***Figure 3***). Furthermore, the promoters of tissue-specific gene sets were analyzed to identify enriched CREs. The PlantPan 4.0 TF binding site tool (http://plantpan.itps.ncku.edu.tw/index.html) was used to map a position weight matrix (PWM) of known plant TF binding motifs to promoters of all genes in the *Sorghum bicolor (cv BTx623)* genome. The putative promoter sequences of all sorghum genes were limited to 1500 bp upstream from −1 position from the start codon of the gene. The enrichment of CREs in the promoter regions of tissue-specific genes were calculated based on frequency of occurrence of CREs in the promoter regions compared to occurrence in all promoters in the genome (fold change ≥ 1.5, adjusted FDR ≤ 0.05). Here, CREs that were present in both (+) and (-) strand were considered. The enriched CREs in each stem-specific gene set (AOSS, TSS-Early, TSS-Late) is visualized in ***Figure 4***.

### Stem-preferred gene regulatory network (GRN) construction

We constructed five gene regulatory networks, in which pairwise correlation analysis (Pearson correlation ≥ |0.7| and Mutual Rank ≥ |0.7|, p-value ≤ 0.05) was performed for all stem-preferred genes (AOSS, TSS-Early, TSS-Late) and stem-expressed TFs accounting for CREs enriched for each stem-preferred gene and stem-expressed TFs (Obayashi and Kinoshita, no date). Gene-gene edges are defined by Pearson Correlation Coefficients (PCC) over 0.7 with a Mutual Rank (MR) > 0.7. The MR is another co-expression measure which takes a geometric average of the PCC rank from gene A to gene B and that of gene B to gene A. The use of MR measure makes sure only highly significant co-expressed gene pairs are considered. TF-target associations were predicted using known plant TF binding motifs mapped to 1.5 kbp upstream from the start of each gene, in which only enriched CREs were considered. The final GRN is a merged network that contain TF-to-gene connections, which have highly correlated expression. Networks were constructed using a custom R script (https://github.com/Madara-Dilhani/GeneReg_Network) and visualized using Cytoscape (version 3.8). Network analysis was performed in NetworkAnalyzer tool in Cytoscape to assess network properties.

### Variation in stem-preferred GRN of Grain Sorghum and Sweet Sorghum genotypes

We used RNA-seq data of *Sorghum bicolor cv Della* stem tissue for 11 different time points covering five development stages from McKinley et al. (2018) to evaluate conserved gene regulation of stem-preferred genes between sweet and grain sorghum varieties. McKinley et al. (2018) had sampled *Sorghum bicolor cv Della* stem tissue samples by taking the 10th internode of the stem at the following time points: Days after emergence (DAE), days before Anthesis (DBA), and days after anthesis (DAA). At the juvenile stage: 14 DAE/-55 DBA; at the vegetative stage: 29 DAE/-40 DBA, 31 DAE/-38 DBA, 40 DAE/-29 DBA; at floral initiation: 53 DAE/ -16 DBA, 62 DAE/-7 DBA; at anthesis: 69 DAE/Anthesis; at grain maturity: 80 DAE/+11 DAA, 94 DAE/+25 DAA; at post-grain maturity: 102 DAE/+43 DAA, 137 DAE/+68 DAA. We used publicly available raw counts for all the samples obtained from Phytozome to normalize gene expression levels using modules from edgeR package (Robinson, McCarthy and Smyth, 2010) in R (R Core Team, 2019). We constructed eleven gene regulatory networks (GRNs) for 11 time points sampled representing 5 developmental stages. The co-expression analysis (Pearson correlation ≥ |0.7| and Mutual Rank ≥ |0.7|, p-value ≤ 0.05) was performed for all stem-preferred genes (AOSS, TSS-Early, TSS-Late) and expressed TFs accounting for CREs enriched for each stem-preferred gene and stem-expressed TFs (Obayashi & Kinoshita, 2009). Gene regulatory networks between BTx623 and Della were compared for comparable time points, which were selected based on time of sample collection in their respective developmental maturities and anatomical similarity (McKinley et al., 2018). The comparable time points were as follows: a) Juvenile - BTx623-8 DAE with Della-14 DAE, b) Vegetative - BTx623-24 DAE with Della-29 DAE, c) Anthesis - BTx623-65 DAE with Della-69 DAE, d) Grain Maturity - BTx623-96 DAE with Della-94 DAE.

### Single nucleotide and structural variations in regulatory regions of stem-preferred genes and their regulatory TFs between *Sorghum bicolor cv BTx623* and *Sorghum bicolor cv Della*

We collected Della re-sequenced genome and variant reports (https://genome.jgi.doe.gov/portal/Sorbicsequencing_9/Sorbicsequencing_9.info.html), where authors used the following variant calling pipeline: a) The Della re-sequenced reads were mapped to the BTx623 reference genome using BWA (Li and Durbin, 2009), b) putative single nucleotide point (SNP) mutation and small indel sites were identified by using samtools (Li *et al.*, 2009), bcftools (Danecek, Schiffels and Durbin, 2016), vcfutils (Danecek *et al.*, 2011), picard (*Picard Tools - By Broad Institute*, no date), snpEff, (Cingolani *et al.*, 2012), c) putative large structural variant sites such as deletions, insertions, inversions, intra-chromosomal translocations, inter-chromosomal translocations using following tools; BreakDancer (Fan *et al.*, 2014), Pindel (Ye *et al.*, 2009). For this study, we predicted significant point mutations and structural variants within 1.5 kb upstream promoter region, gene body, and 500 bp downstream region of tissue-specific genes and their regulators in respective developmental stages from BTx623 gene regulatory network to identify variable and conserved regulatory regions.

### Data availability statement

There is no data associated with this manuscript. R script for tissue-specificity analysis is available at https://github.com/Madara-Dilhani/Tissue_Specificity. Other scripts used for network analysis are available at https://github.com/Madara-Dilhani/GeneReg_Network.

## Acknowledgements

*This work was funded by the DOE Center for Advanced Bioenergy and Bioproducts Innovation (U.S. Department of Energy, Office of Science, Office of Biological and Environmental Research under Award Number DE-SC0018420) and in part by the DOE Great Lakes Bioenergy Research Center (DOE BER Office of Science DE-FC02-07ER64494). Any opinions, findings, and conclusions or recommendations expressed in this publication are those of the author(s) and do not necessarily reflect the views of the U.S. Department of Energy.*

## Author contributions

MHA and AMC conceived the project; MHA, AMC, and JEM designed the experiments; JEM contributed the data; MHA performed all data analysis; MHA, AMC, and JEM wrote the manuscript. All authors reviewed and approved of the manuscript.

## Supporting Information

**Figure S1:** Expression dynamics of highly expressed most stem-specific AOSS genes in Sorghum cv BTx623 and cv Della.

**Figure S2:** The view upstream non-coding region, gene body, downstream non-coding region of 3 highly expressed most stem-specific AOSS genes from Integrated Genomic Viewer (IGV).

**Figure S3:** The view upstream non-coding region, gene body, downstream non-coding region of AOSS genes with significant genetic alterations from Integrated Genomic Viewer (IGV).

**Figure S4:** The Integrated Genomic Viewer (IGV) view upstream non-coding region, gene body, downstream non-coding region of regulatory TFs of Sobic.006G024800 during a) Early stage and b) Late stage.

**Table S1:** Previously known xylem and phloem specific cis-regulatory element in different vascular specific promoters

**Table S2:** Statistics of tissue-specific genes across tissues during development **Table S3:** Tissue-specificity Index by stages for AOSS, TSS-Early, TSS-Late genes **Table S4:** Transcription factors which are TF hubs in at least one of the 15 GRNs

**Table S5:** Transcription factors which showed change of TF hub characteristics in early and late developmental stages

**File S1:** The list of predicted single nucleotide variations (SNVs) between BTx623 and Della found in the gene body, 1.5 kbp upstream (including predicted CREs) for stem-expressed genes in the AOSS, TSSE, and TSSL gene group and their TF regulators.

**File S2:** The list of predicted structural variations (SVs) between BTx623 and Della found in the gene body, 1.5 kbp upstream (including predicted CREs) for stem-expressed genes in the AOSS, TSSE, and TSSL gene group and their TF regulators.

**Data S1:** TF–target gene interactions for each of 15 GRNs generated for this study is presented in.sif files.

