## Supplemental Figures for "*Sorghum bicolor* cultivars have divergent and dynamic gene regulatory networks that control the temporal expression of genes in stem tissue"

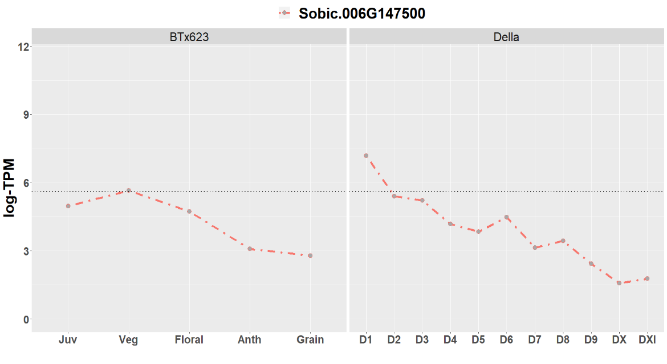

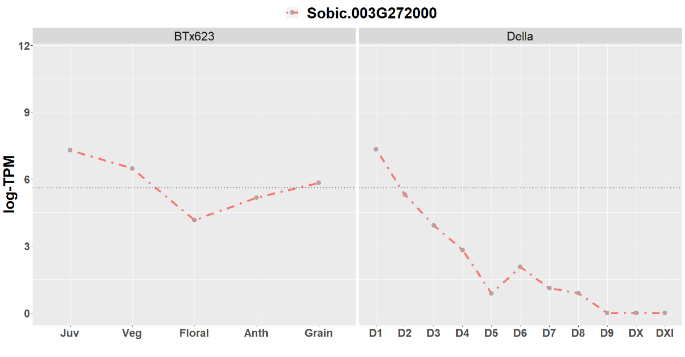

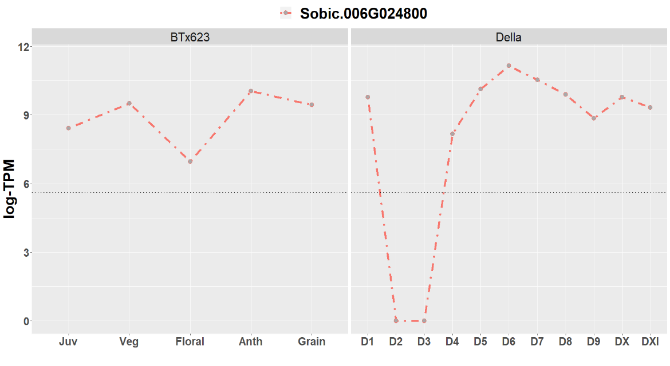

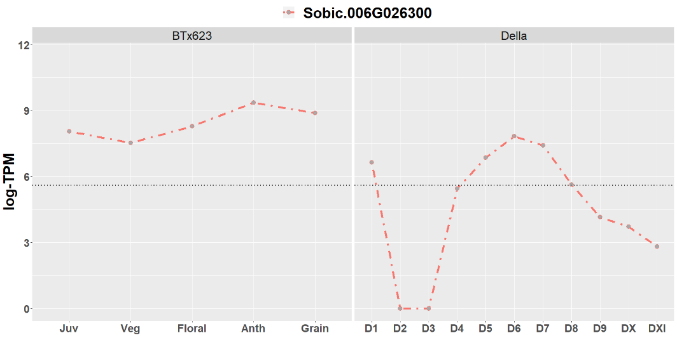

**Figure S1: Expression dynamics of highly expressed most stem-specific AOSS genes in Sorghum cv BTx623 and cv Della.**

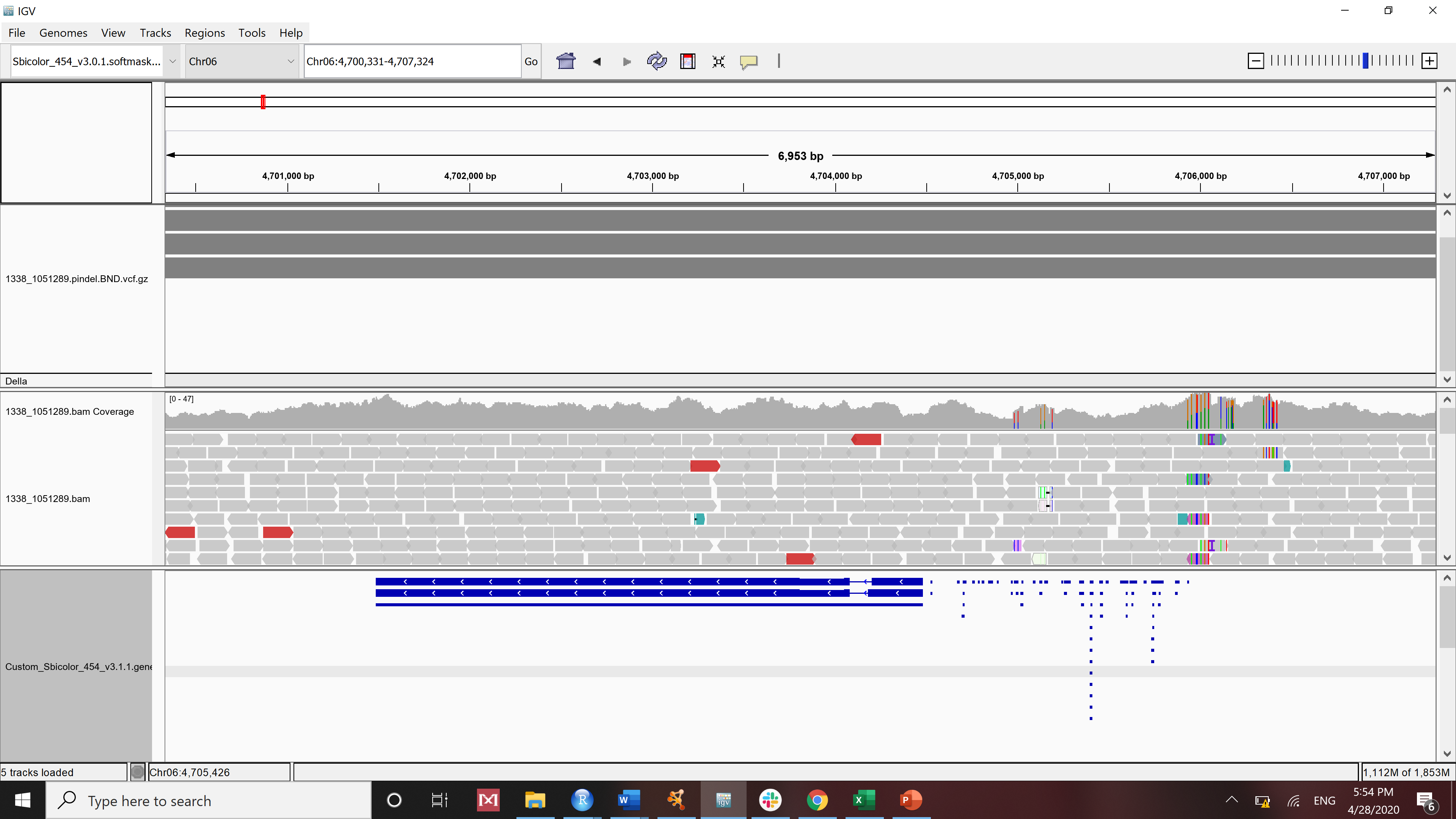

Sobic.006G026300

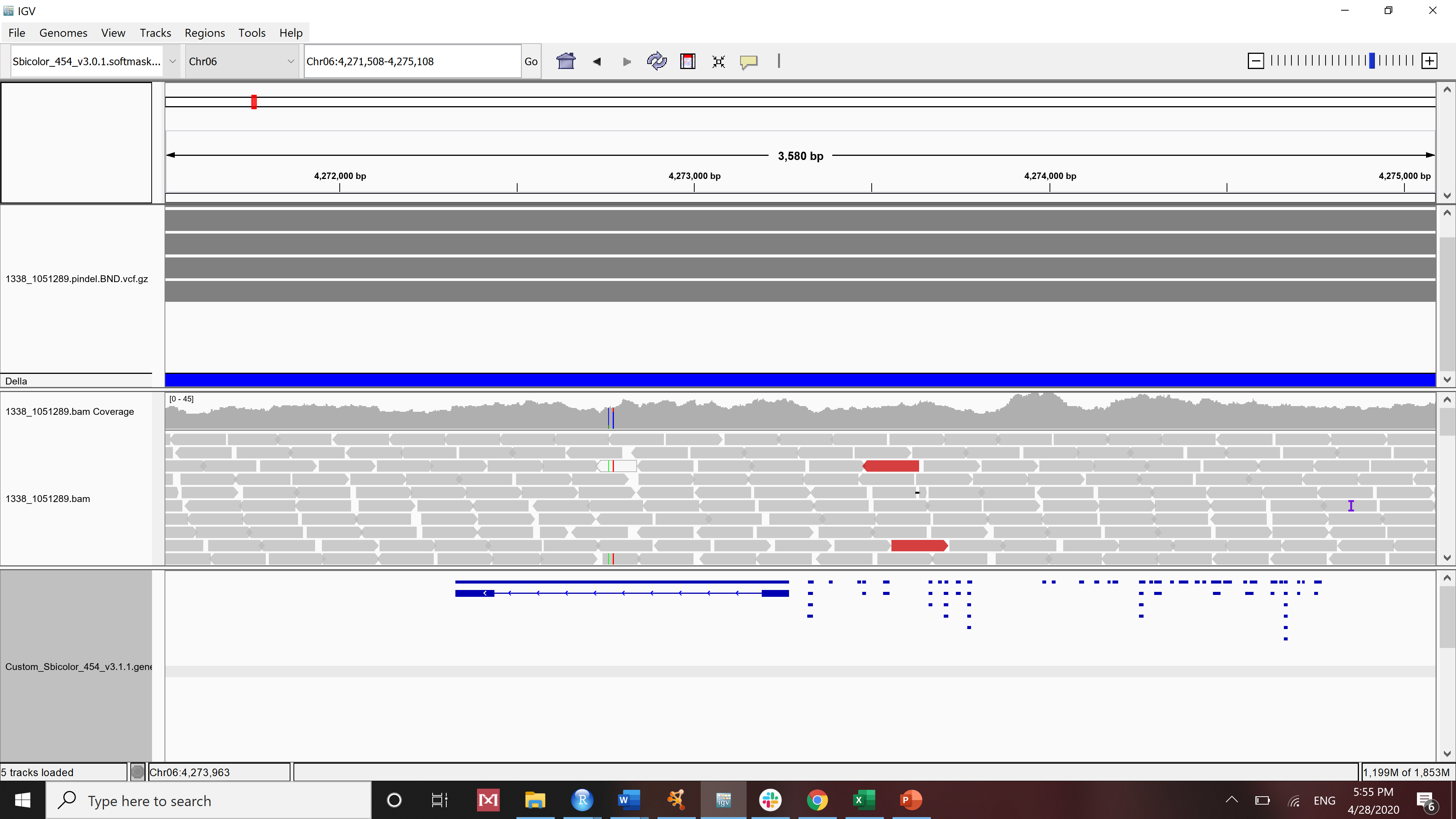

Sobic.006G024800

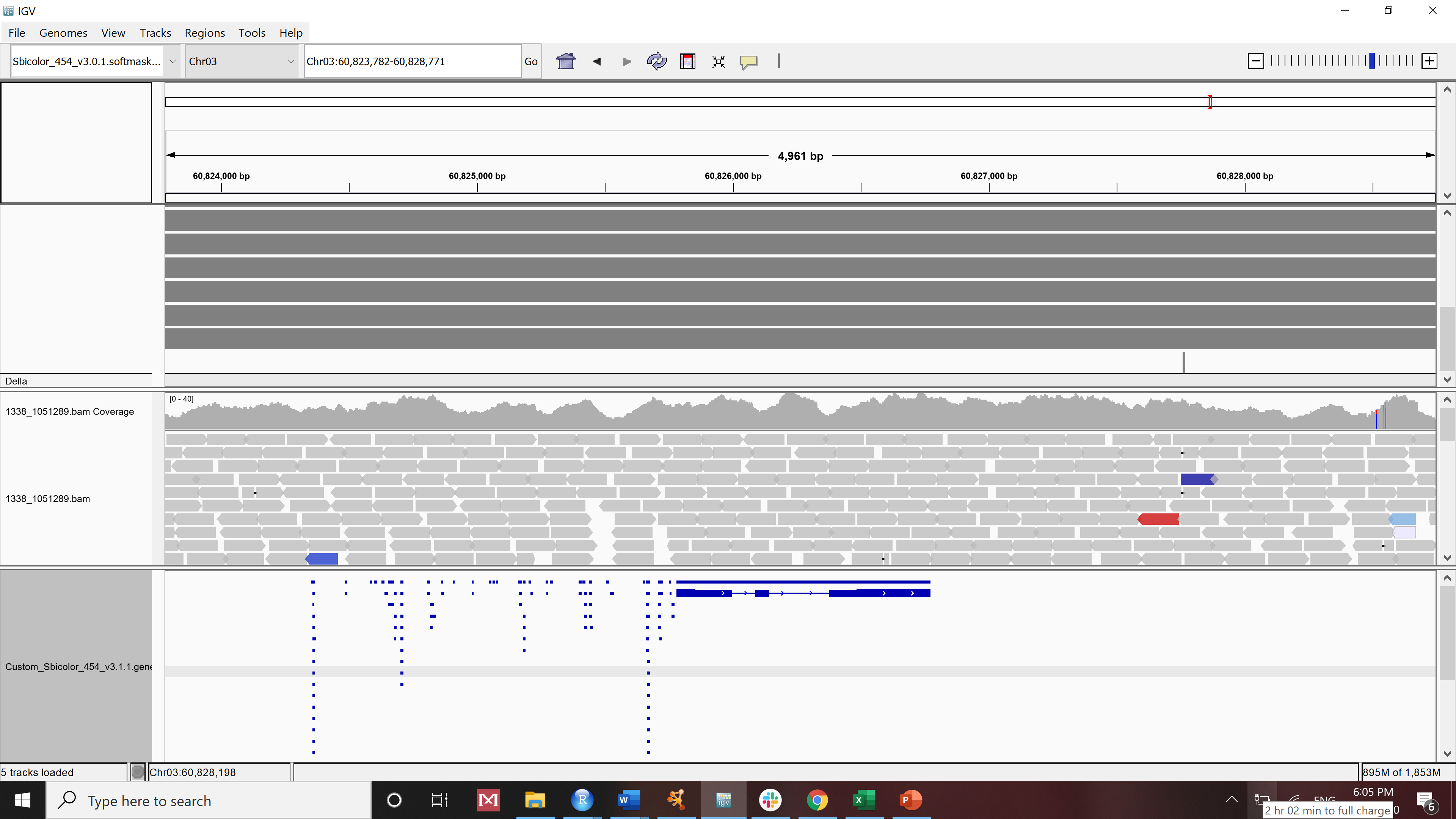

Sobic.003G272000

**Figure S2: The view upstream non-coding region, gene body, downstream non-coding region of 3 highly expressed most stem-specific AOSS genes from Integrated Genomic Viewer (IGV).** Histogram in the top of each IGV window shows the sum of the aligned sequencing reads along the genome (read coverage). Genes are represented as lines and boxes. Lines represent intronic regions, and boxes represent exonic regions. The arrows indicate the direction/strand of transcription for the gene. Identified enriched CRE elements in our study is visualized in individual squares upstream of the gene.

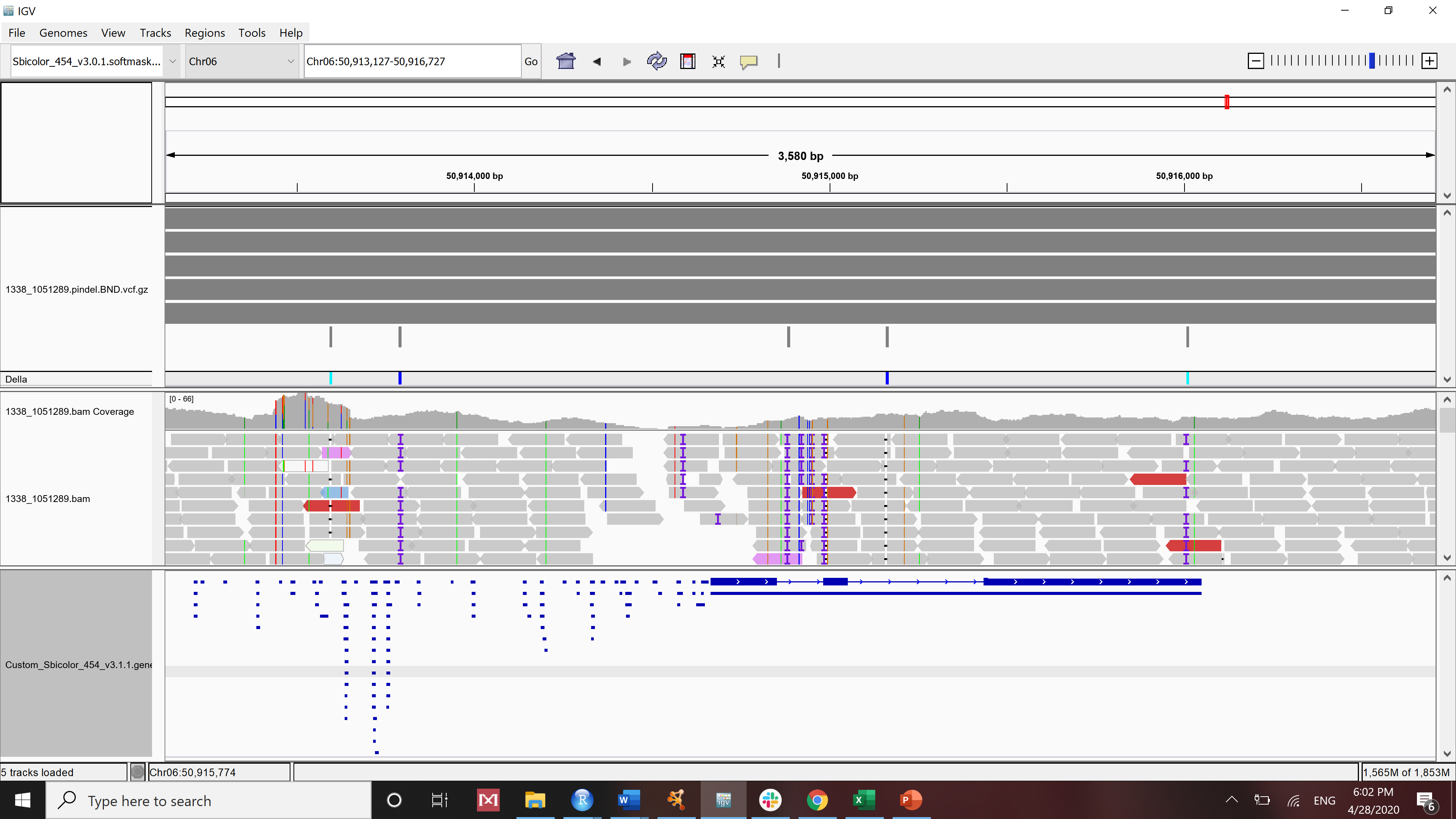

**Sobic.006G147500 (LSH6)**

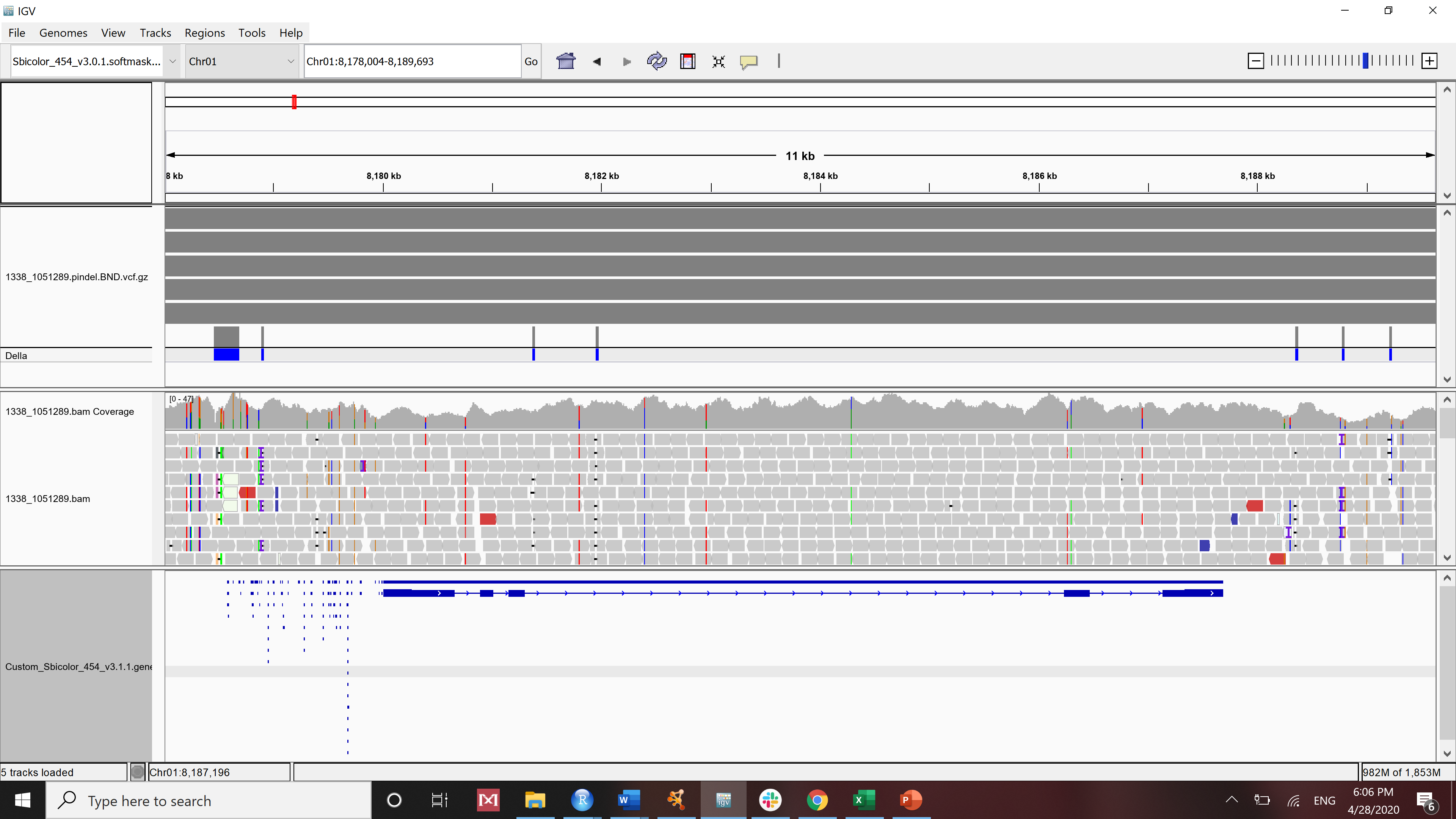

**Sobic.001G106200 (KNAT1)**

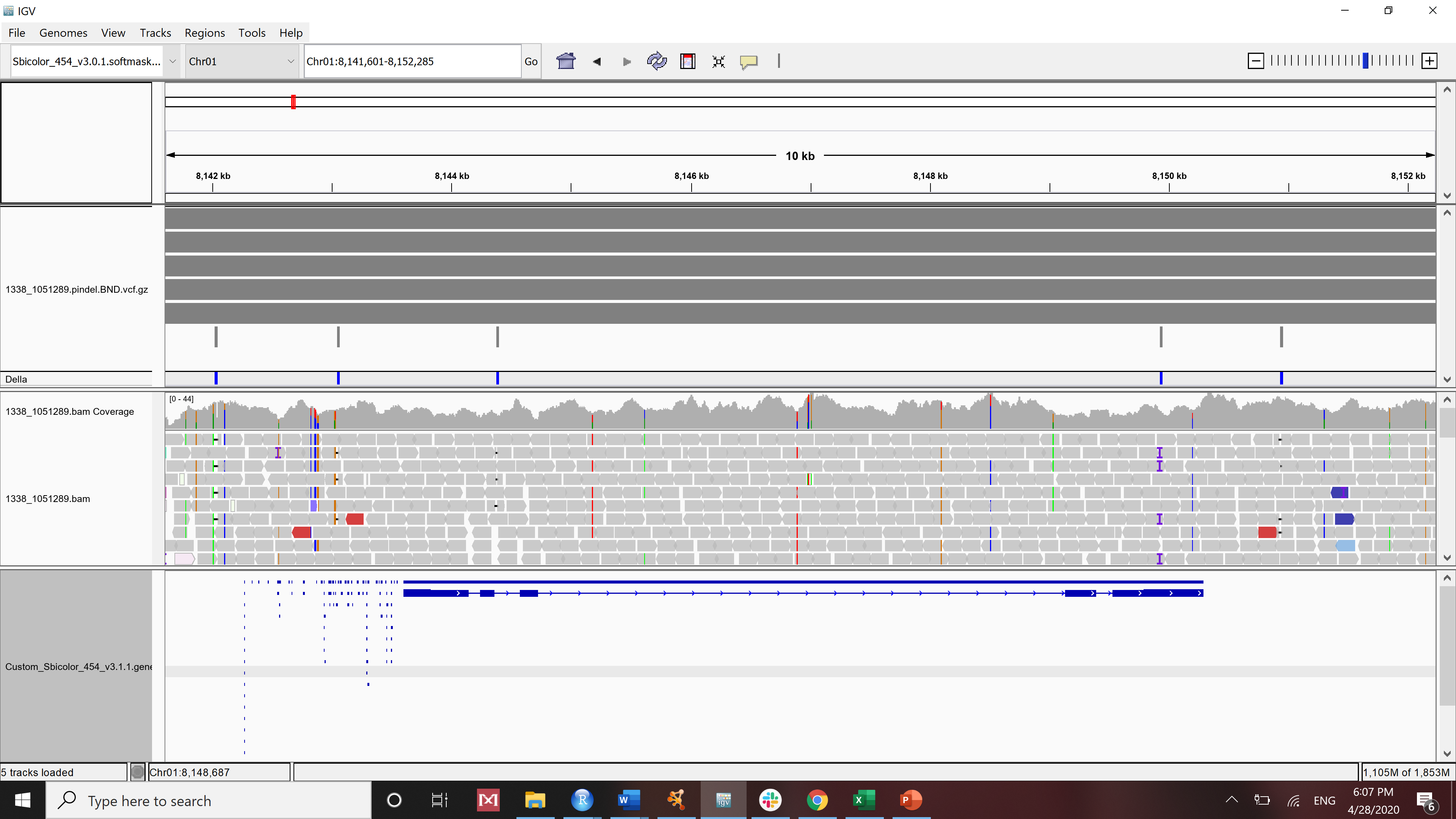

**Sobic.001G106000 (KNAT1)**

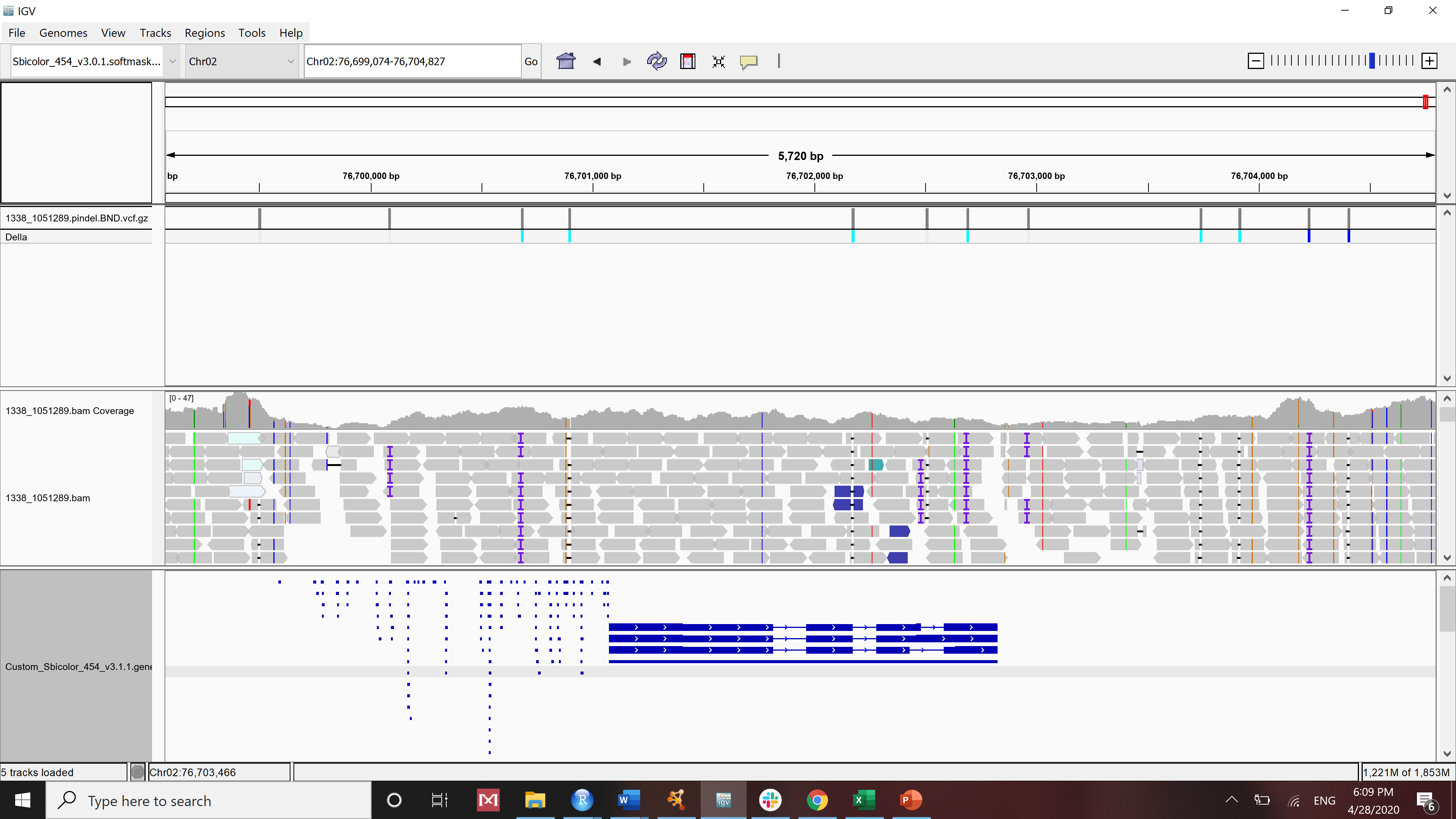

**Sobic.002G419500 (HAD)**

**Figure S3: The view upstream non-coding region, gene body, downstream non-coding region of AOSS genes with significant genetic alterations from Integrated Genomic Viewer (IGV).** Histogram in the top of each IGV window shows the sum of the aligned sequencing reads along the genome (read coverage). Genes are represented as lines and boxes. Lines represent intronic regions, and boxes represent exonic regions. The arrows indicate the direction/strand of transcription for the gene. Identified enriched CRE elements in our study is visualized in individual squares upstream of the gene.

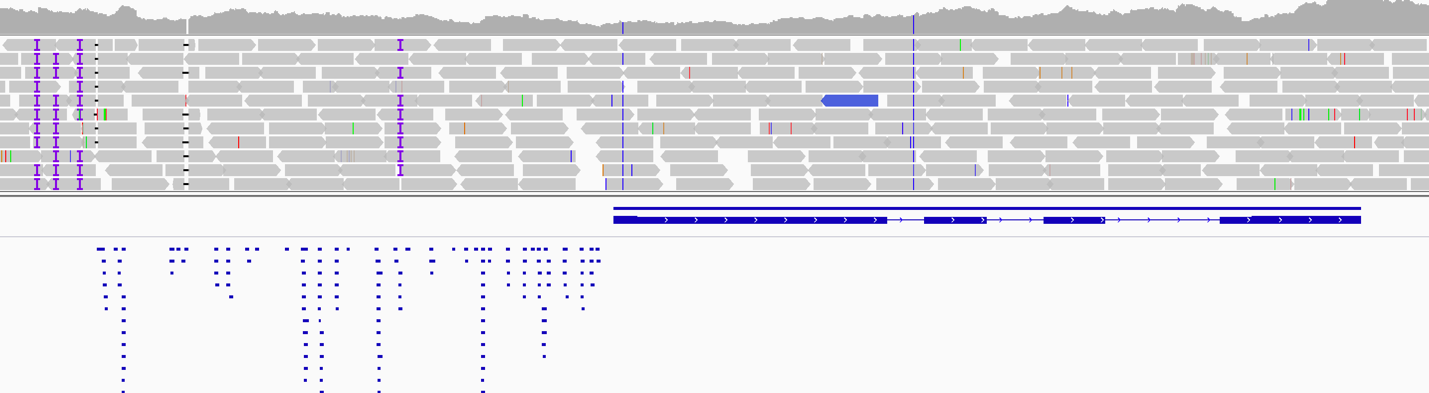

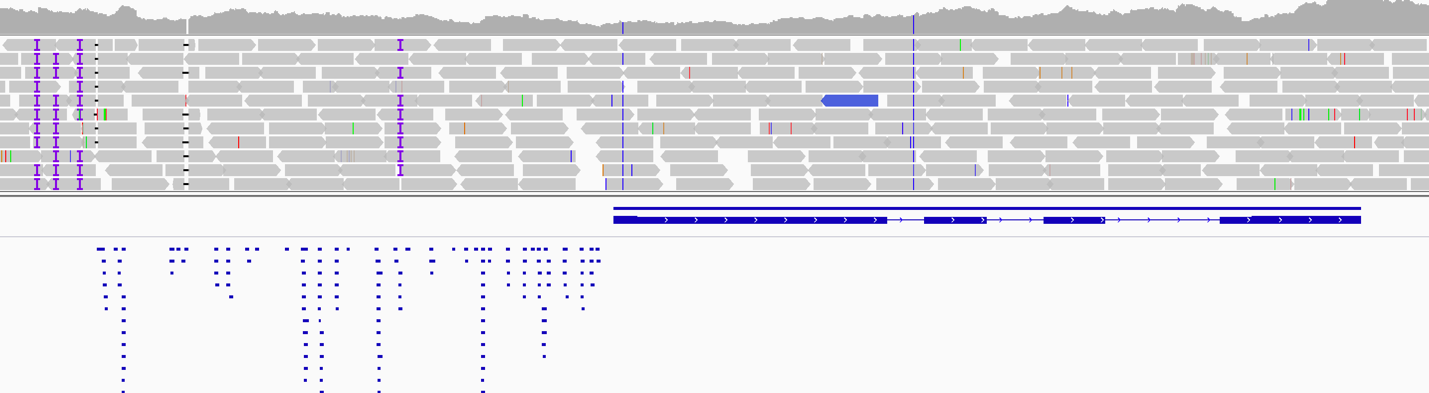

**Sobic.001G513700**

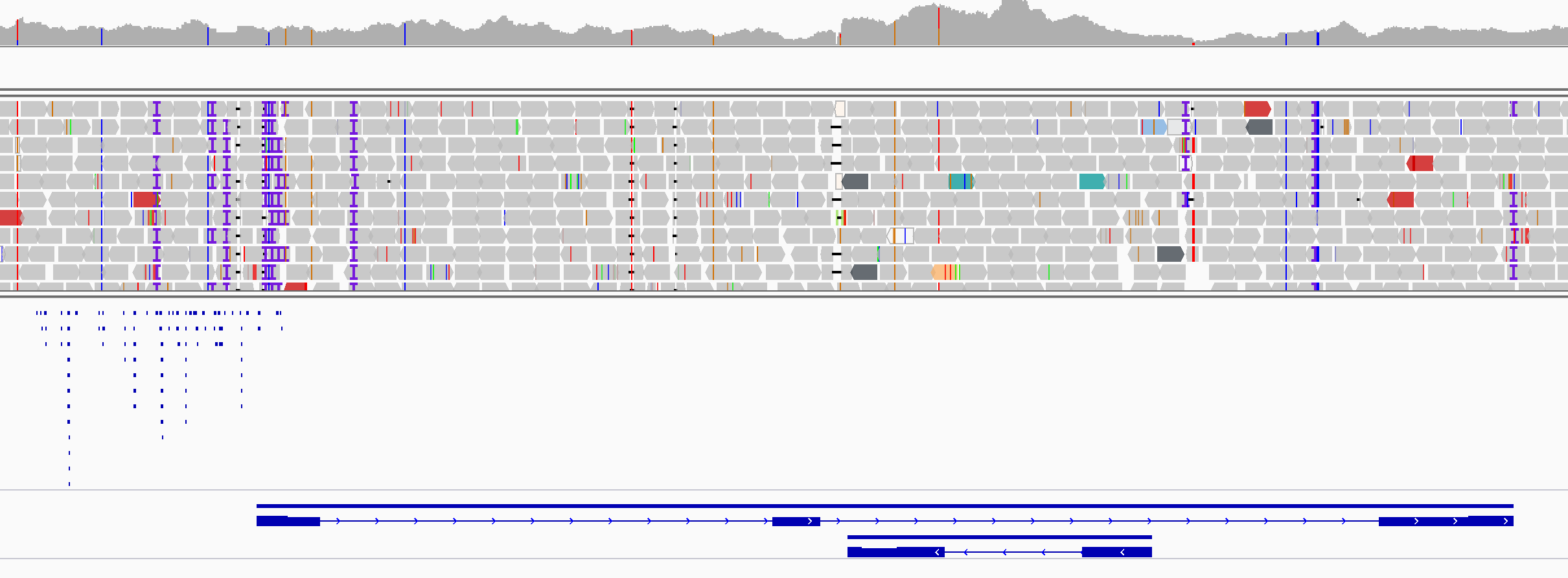

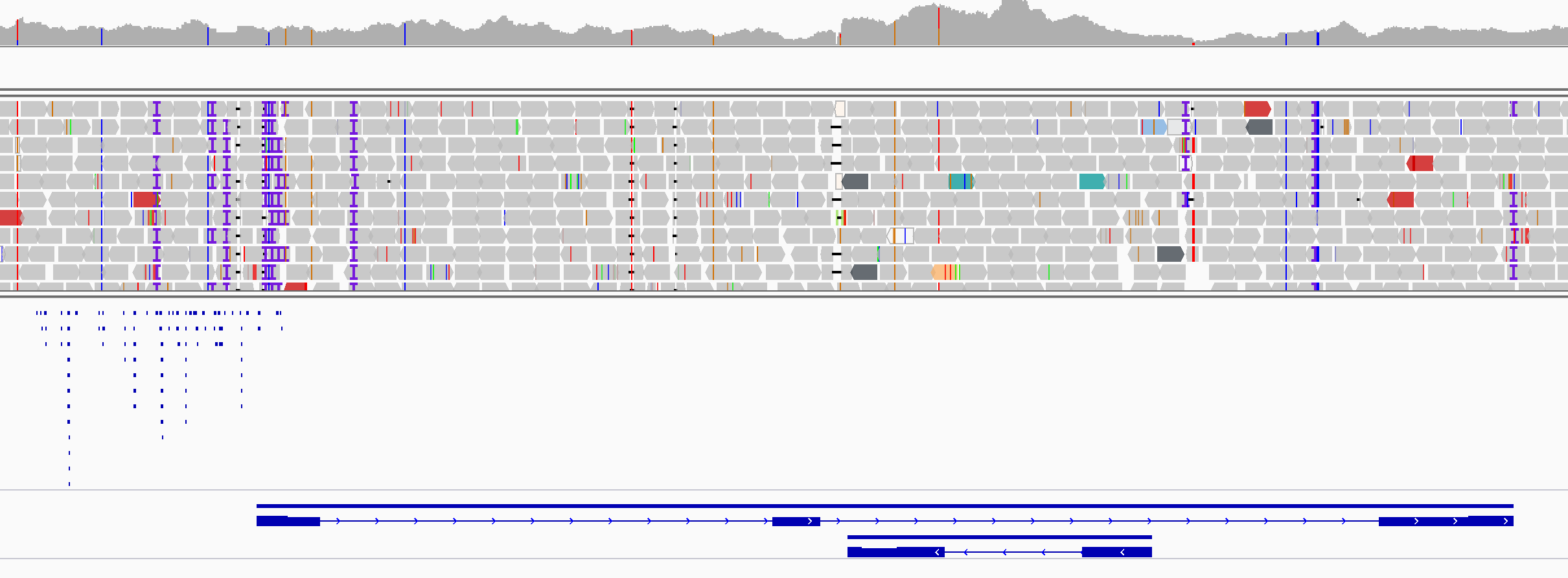

**Sobic.008G164800**

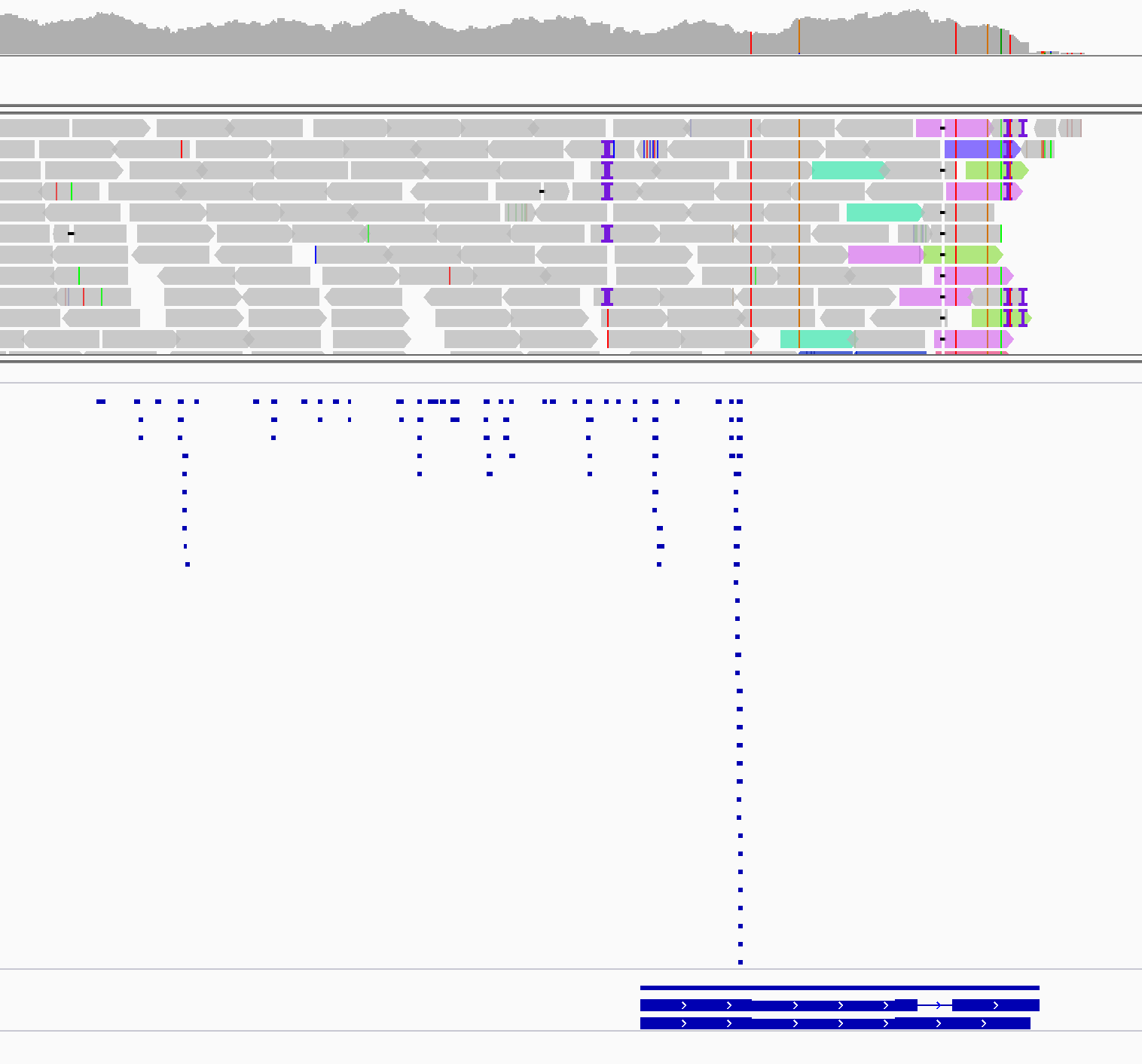

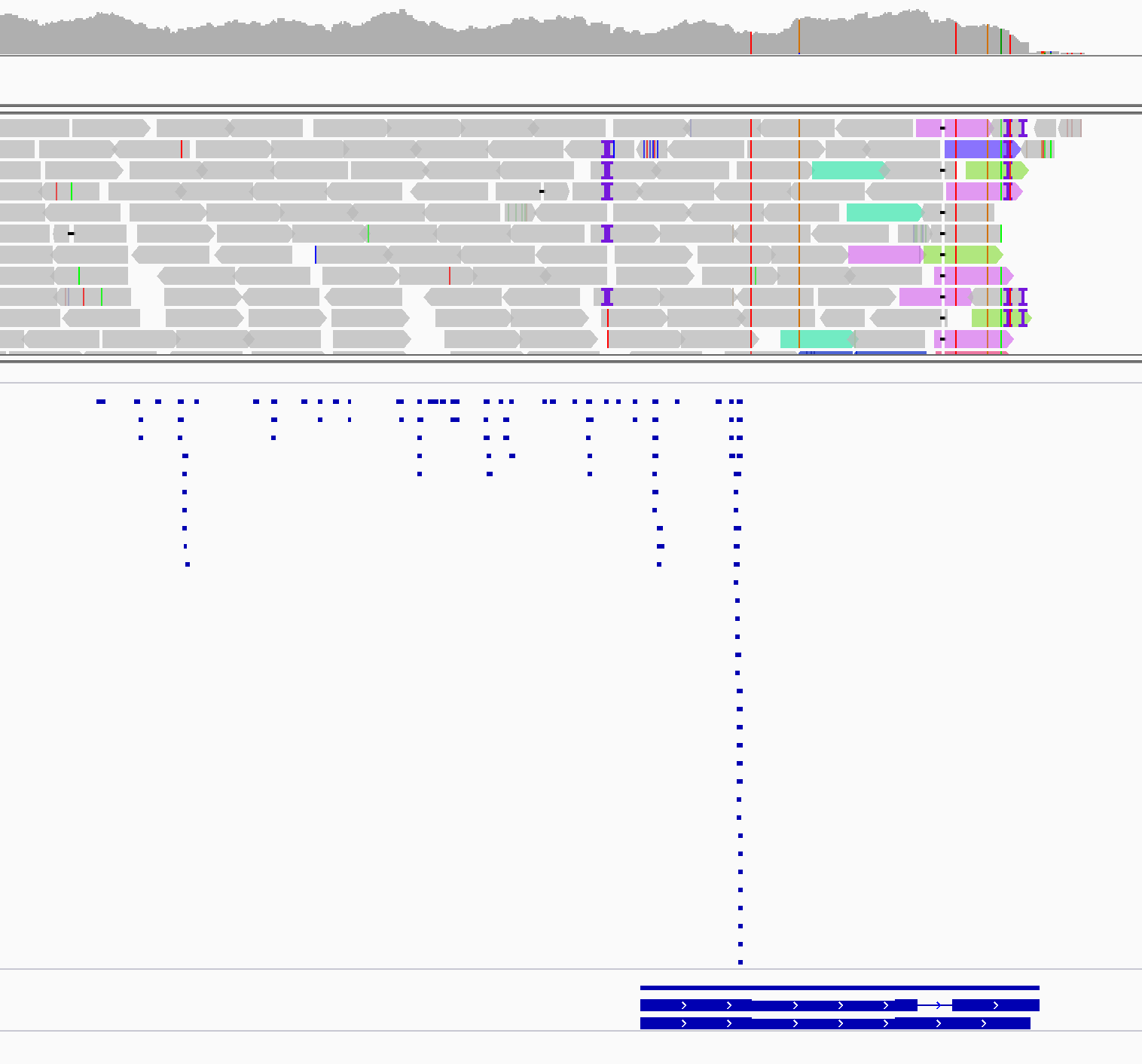

**Sobic.005G019800**

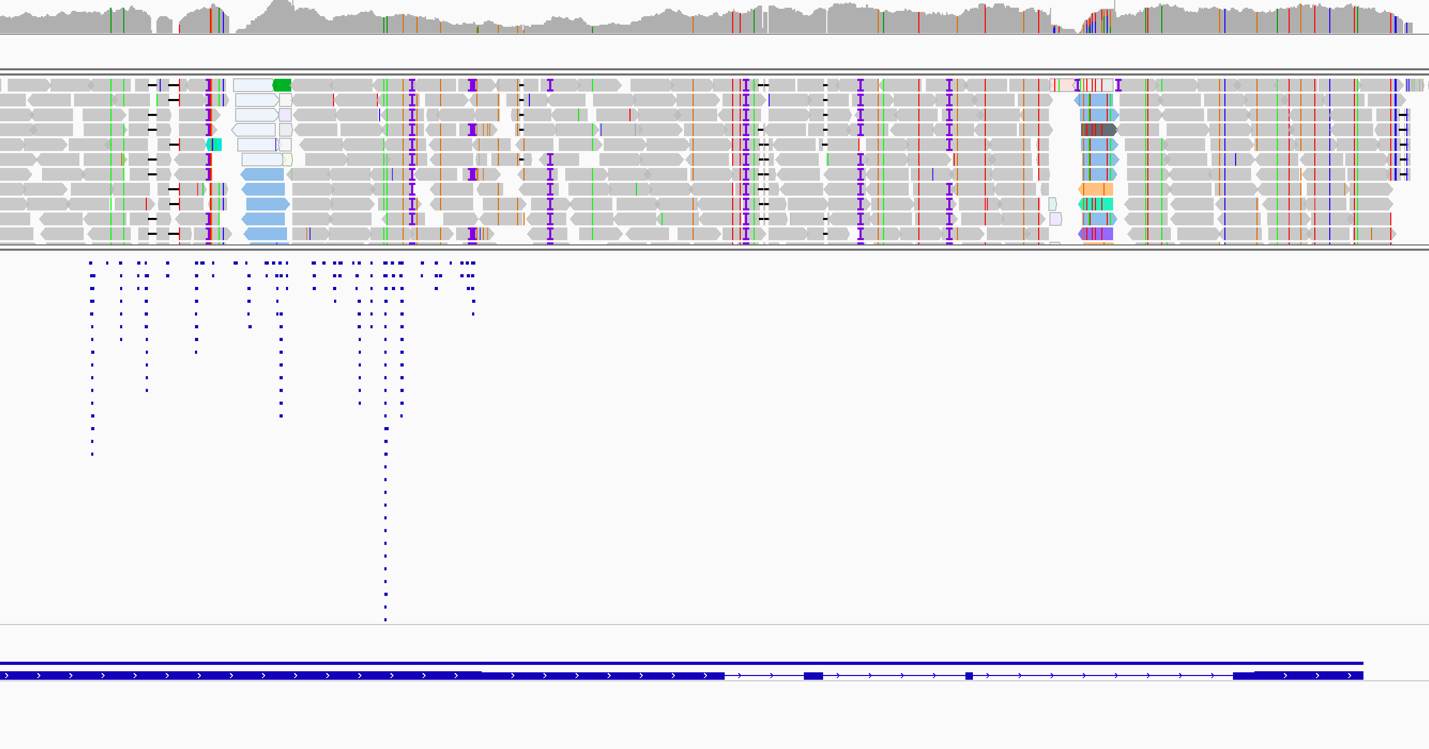

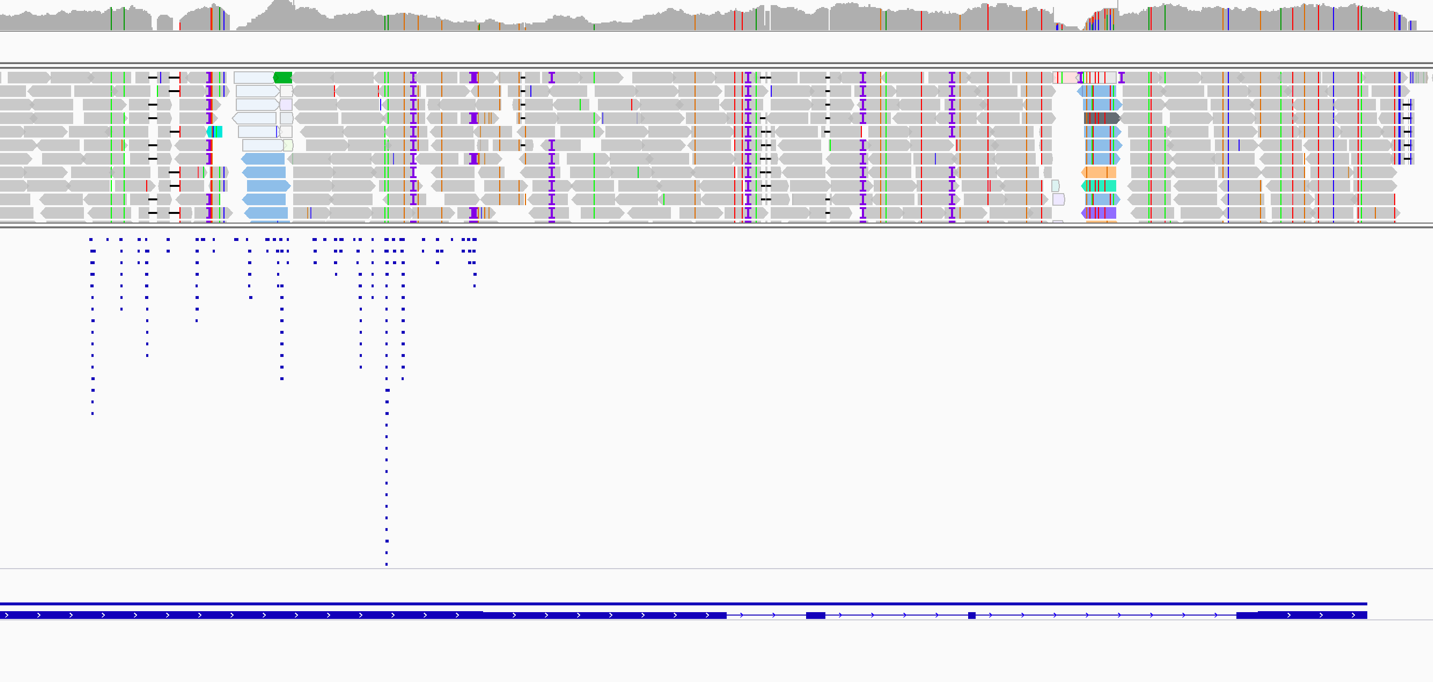

**Sobic.002G225100**

**Figure S4: The Integrated Genomic Viewer (IGV) view upstream non-coding region, gene body, downstream non-coding region of regulatory TFs of Sobic.006G024800 during a) Early stage and b) Late stage.** Histogram in the top of each IGV window shows the sum of the aligned sequencing reads along the genome (read coverage). Genes are represented as lines and boxes. Lines represent intronic regions, and boxes represent exonic regions. The arrows indicate the direction/strand of transcription for the gene. Identified enriched CRE elements in our study is visualized in individual squares upstream of the gene.
